# Genetic and environmental manipulation of *Arabidopsis* hybridization barriers uncover antagonistic functions in endosperm cellularization

**DOI:** 10.1101/2023.05.26.542444

**Authors:** Katrine N. Bjerkan, Renate M. Alling, Ida V. Myking, Anne K. Brysting, Paul E. Grini

**Affiliations:** Section for Genetics and Evolutionary Biology, University of Oslo, 0316 Oslo, Norway; CEES, Department of Biosciences, University of Oslo, 0316 Oslo, Norway

**Keywords:** *Arabidopsis thaliana*, *A. arenosa*, *A. lyrata*, hybrid barrier, temperature, endosperm, seed development

## Abstract

Speciation by reproductive isolation can occur by hybridization barriers acting in the endosperm of the developing seed. The nuclear endosperm is a nutrient sink, accumulating sugars from surrounding tissues, and undergoes coordinated cellularization, switching to serve as a nutrient source for the developing embryo. Tight regulation of cellularization is therefore vital for seed and embryonic development. Here we show that hybrid seeds from crosses between *Arabidopsis thaliana* as maternal contributor and *A. arenosa* or *A. lyrata* as pollen donors result in an endosperm based post-zygotic hybridization barrier that gives rise to a reduced seed germination rate. Hybrid seeds display opposite endosperm cellularization phenotypes, with late cellularization in crosses with *A. arenosa* and early cellularization in crosses with *A. lyrata*. Stage specific endosperm reporters display temporally ectopic expression in developing hybrid endosperm, in accordance with the early and late cellularization phenotypes, confirming a disturbance of the source-sink endosperm phase change. We demonstrate that the hybrid barrier is under the influence of abiotic factors, and show that a temperature gradient leads to diametrically opposed cellularization phenotype responses in hybrid endosperm with *A. arenosa* or *A. lyrata* as pollen donors. Furthermore, different *A. thaliana* accession genotypes also enhance or diminish seed viability in the two hybrid cross-types, emphasizing that both genetic and environmental cues control the hybridization barrier. We have identified an *A. thaliana* MADS-BOX type I family single locus that is required for diametrically opposed cellularization phenotype responses in hybrid endosperm. Loss of AGAMOUS-LIKE 35 significantly affects the germination rate of hybrid seeds in opposite directions when transmitted through the *A. thaliana* endosperm, and is suggested to be a locus that promotes cellularization as part of an endosperm based mechanism involved in post-zygotic hybrid barriers. The role of temperature in hybrid speciation and the identification of distinct loci in control of hybrid failure have great potential to aid the introduction of advantageous traits in breeding research and to support models to predict hybrid admixture in a changing global climate.

## 1 Introduction

Speciation is usually a continuous process towards increasing divergence and reproductive isolation between two lineages. Reproductive isolation can be obtained due to hybridization barriers, which act before fertilization (pre-zygotic) or after fertilization (post-zygotic) (Rieseberg and Willis, 2007; Widmer et al., 2009). A special case of post-zygotic hybridization barriers acts in the developing seed (Lafon-Placette and Köhler, 2016), resulting in developmental abnormality in the endosperm (Cooper and Brink, 1942). Seed death is accepted to be mainly due to failure of endosperm development since the embryo can be rescued in culture after microdissection (Sharma, 1999). Many studies have indicated endosperm deficiency to be the major cause of hybrid seed inviability (Brink and Cooper, 1947; Sukno et al., 1999; Dinu et al., 2005; Roy et al., 2011) and an endosperm-based hybridization barrier has been shown to be conserved across distinct species groups such as *Arabidopsis* (Lafon-Placette and Köhler, 2016), *Capsella* (Rebernig et al., 2015; Dziasek et al., 2021), rice (Ishikawa et al., 2011; Sekine et al., 2013; Zhang et al., 2016; Tonosaki et al., 2018; Wang et al., 2018), tomato (Florez-Rueda et al., 2016; Roth et al., 2019), monkeyflower (Oneal et al., 2016; Flores-Vergara et al., 2020; Kinser et al., 2021), and potato (Johnston and Hanneman, 1982; Cornejo et al., 2012). This suggests that the phenomenon represents a major mechanism of reproductive isolation in plants, however it is largely ignored in modern literature of speciation (Lafon-Placette and Köhler, 2016).

The endosperm is a triploid tissue that requires tight control of genome dosage (2:1 maternal:paternal ratio). Cellularization of the endosperm marks a transition in seed development, as up to this point, the endosperm functions as a nutrient sink. At this developmental time point, the endosperm concurrently switches from nutrient sink to source for the developing embryo (Lafon-Placette and Köhler, 2016), and manipulating the timing of endosperm cellularization through interploidy crosses arrests embryo development (Scott et al., 1998; Hehenberger et al., 2012). Similarly, hybridization between plant species was shown to result in embryo arrest due to endosperm cellularization failure (Haig and Westoby, 1988, 1991; Comai et al., 2000; Bushell et al., 2003). Manipulating the ploidy of parents in interspecies crosses has also shown to improve the success of hybridization, demonstrating a requirement for genome balance in the endosperm (Comai et al., 2000; Bushell et al., 2003; Lafon-Placette et al., 2017). Genomic imprinting is an epigenetic phenomenon, which infers parent-of-origin allele specific expression of maternally or paternally inherited alleles (Hornslien et al., 2019; Batista and Kohler, 2020). As proper endosperm development depends on a correct ratio of parental genomes, it is suggested that differences in genomic imprinting programs may be responsible for the evolution of sexual incompatibility in crosses between divergent individuals (Haig and Westoby, 1988, 1991; Bushell et al., 2003; Schatlowski and Kohler, 2012). Alternatively, or additionally, epigenetic remodeling upon hybridization due to combination of diverged maternal and paternal siRNAs may lead to comprehensive failure of genomic imprinting and ectopic expression of transposons and imprinted genes (Martienssen, 2010; Ng et al., 2012). Recent evidence supports this emerging role of imprinted genes, and in *Arabidopsis* interspecies hybrids, paternally expressed genes (PEGs) shifted from paternally biased to be maternally expressed (MEGs) (Josefsson et al., 2006). Importantly, both PEGs and MEGs have been shown to erect hybridization barriers, and mutational loss of these genes has been reported to bypass hybridization barriers in interspecies crosses (Walia et al., 2009; Wolff et al., 2015). A major part of the MEGs and PEGs encodes proteins that activate pathways in the endosperm consistent with the prominent role of cellularization in seed survival.

Proper endosperm development in *Arabidopsis* is reliant on the FERTILIZATION INDEPENDENT SEED-Polycomb Repressive Complex 2 (FIS-PRC2), which is a H3K27-methyltransferase primarily composed of FIS2, MEDEA (MEA), FERTILIZATION-INDEPENDENT ENDOSPERM (FIE) and MULTICOPY SUPPRESSOR of IRA1 (MSI1) (Grossniklaus et al., 1998; Kiyosue et al., 1999; Luo et al., 1999; Köhler et al., 2003). FIS-PRC2 is important in endosperm development indirectly through the genes it regulates, which include several type I MADS-box transcription factors (TFs) (Zhang et al., 2018). Deregulation of type I MADS-box TFs in interspecies crosses has been postulated to induce the endosperm-based hybridization barrier, but unfortunately most of these TFs have no clear function because of extensive genetic redundancy (De Bodt et al., 2003; ParLenicová et al., 2003; Walia et al., 2009; Bemer et al., 2010). One exception is *AGAMOUS-LIKE 62* (*AGL62*), which is found to suppress cellularization of the endosperm in *Arabidopsis* (Kang et al., 2008). The clear function of AGL62 is further emphasized through its interaction with the FIS-PRC2 complex (Hehenberger et al., 2012), with mutation of FIE, MEA, FIS2 and MSI1 resulting in an ectopic proliferation of nuclear endosperm (Grossniklaus et al., 1998; Köhler et al., 2003; Guitton et al., 2004), whereas the *agl62* mutant results in precocious cellularization (Kang et al., 2008). AGL62 mutation has also been shown to alleviate the hybridization barrier in the *A. thaliana* × *A. arenosa* cross, resulting in a higher germination rate (Walia et al., 2009; Bjerkan et al., 2020).

Recent advances in elucidating molecular mechanisms and genetic networks in hybrid endosperm lethality suggest that imprinted genes and genetic variation in the hybrid parents are important factors that can enhance or repress the frequency of the endosperm-based barrier (Bushell et al., 2003; Josefsson et al., 2006; Walia et al., 2009; Burkart-Waco et al., 2012, 2013, 2015; Kradolfer et al., 2013; Schatlowski et al., 2014; Rebernig et al., 2015; Wolff et al., 2015; Bjerkan et al., 2020). However, the mechanistic role of these factors and the interplay of genetic networks is largely unknown. For instance, ploidy can bypass the endosperm-based post-zygotic barrier, but a general role of ploidy cannot be attributed since ploidy plays a different role in maternal and paternal cross settings (Lafon-Placette et al., 2017). Furthermore, the role of gene dosage and genomic imprinting is supported by reports suggesting that mutation of MEGs and PEGs can overcome the endosperm-based post-zygotic barrier (Dilkes et al., 2008; Walia et al., 2009; Wolff et al., 2015; Borges et al., 2018), but a general role of imprinted genes cannot be defined since only some imprinted genes appear to have this effect (Walia et al., 2009; Burkart-Waco et al., 2015; Rebernig et al., 2015; Wolff et al., 2015). Since different accessions of the parental individuals in a hybrid cross can partly bypass the endosperm-based post-zygotic barrier without any change in ploidy (Walia et al., 2009; Wolff et al., 2015), the role of maternal and paternal genomes cannot be generalized.

The genus *Arabidopsis* has been widely used for studying evolutionary questions (Koenig and Weigel, 2015; Koch, 2019) including the effects of interspecific hybridization in controlled crossings (Chen et al., 1998; Comai et al., 2000; Nasrallah et al., 2000; Josefsson et al., 2006; Walia et al., 2009; Burkart-Waco et al., 2012, 2013, 2015; Bjerkan et al., 2020). When diploid *A. thaliana* is crossed to diploid *A. arenosa*, the endosperm shows late cellularization and high degree of seed abortion (Josefsson et al., 2006; Bjerkan et al., 2020). Furthermore, when *A. arenosa* is crossed as male to *A. lyrata*, the same phenotype can be seen with late endosperm cellularization and a very high seed lethality (Lafon-Placette et al., 2017). Interestingly, crossing *A. lyrata* as male to *A. arenosa* results in the opposite effect with early endosperm cellularization (Lafon-Placette et al., 2017).

Here we report that hybrid seeds from crosses between *A. thaliana* mothers with *A. arenosa* or *A. lyrata* pollen donors result in diametrically opposed endosperm phenotypes, both giving rise to reduced seed germination rate, albeit caused by late cellularization in crosses with *A. arenosa* and early cellularization in crosses with *A. lyrata*. We demonstrate that the hybrid barriers are under the influence of abiotic factors, and show that a temperature gradient leads to opposed cellularization phenotype and seed viability in hybrid endosperm with *A. arenosa* or *A. lyrata* as pollen donors. In addition, *A. thaliana* accession genotypes also influence seed viability in the two hybrid cross-types in opposite directions. Using stage specific endosperm reporters, we demonstrate that the source-sink endosperm phase change is delayed or precocious in seeds of the two hybrids. Our data suggests an *A. thaliana* type I MADS-BOX family locus to act as a promoter of endosperm cellularization, affecting the germination rates of *A. arenosa* or *A. lyrata* hybrid seeds in opposite directions.

## 2 Materials and Methods

### 2.1 Plant material and growth conditions

*A. thaliana* accessions (Col-0, C24, Ws-2 and Wa-1) and mutant lines were obtained from the Nottingham Arabidopsis Stock Center (NASC). The *A. arenosa* population MJ09-4 and the *A. lyrata* subsp. *petraea* population MJ09-11 originate from central Europe (Jørgensen et al., 2011; Lafon-Placette et al., 2017; Bjerkan et al., 2020). Mutant lines *agl35-1* (SALK_033801), *agl40-1* (SALK_107011) and *mea-9* (SAIL_724_E07) were in Col-0 accession background (Shirzadi et al., 2011; Kirkbride et al., 2019; Bjerkan et al., 2020). The *agl35-1* T-DNA line was genotyped using primers AAACCAAAGTTTTGCCACTAAGAC, ATTTTTCAGTCAAGATTACCCACC and GCGTGGACCGCTTGCTGCAACTCTCTCAGG. Marker lines *proAT5G09370>>H2A-GFP* (EE-GFP) and proAT4G00220>>H2A-GFP (TE1-GFP) were in Col-0 accession background (van Ekelenburg et al., 2023).

Surface-sterilization of seeds was performed by treatment with 70% ethanol, bleach (20% Chlorine, 0.1% Tween20) and wash solution (0.001% Tween20) for 5 min each step (Lindsey et al., 2017). Sterilized seeds were transferred to petri dishes containing 0.5 MS growth-medium with 2% sucrose (Murashige and Skoog, 1962) and stratified at 4°C for 2 days (*A. thaliana*) or 10 days (*A. lyrata*, *A. arenosa* and hybrids) before germination at 22°C with a 16h/8h light/dark cycle. After two weeks, seedlings were transferred to soil and cultivated at 18°C (16h/8h light/dark cycle, 160 µmol/m^2^/sec, 60-65% humidity). *A. arenosa* strain MJ09-4 was previously demonstrated to be diploid (Bjerkan et al., 2020). *A. lyrata* and *A. thaliana* × *A. lyrata* individual hybrid plants were confirmed diploid by flow cytometry (Supplementary Datasheet S1).

### 2.2 Crosses, temperature- and germination assays

*A. thaliana* plants were emasculated 2 days before pollination. Crossed plants were placed at experimental growth temperature until silique maturity/harvesting. For each cross combination, 15-85 siliques (biological replicates) were harvested individually (Supplementary Datasheet S2). After short-term storage at 4°C seeds were surface-sterilized ON in 5% sodium hypochlorite solution. Seeds from individual siliques were counted and planted individually on 0.5 MS growth-medium and scored for germination by counting protrusions through the seed coat after 10 days at 22°C growth conditions. At day 20, germinated seedlings were checked for *A. thaliana* accidental self-pollination (formation of floral shoots without vernalization).

### 2.3 Microscopy

Feulgen stained seeds were harvested at 6 days after pollination (DAP), stained using Schiff’s reagent (Sigma-Aldrich S5133), fixed and embedded in LR White (London Resin) (Braselton et al., 1996). Imaging was performed using an Andor DragonFly spinning disc confocal microscope with a Zyla4.2 sCMOS 2048×2048 camera attachment and excitation 488 nm / emission 500 to 600 nm. Seeds were scored for embryo and endosperm developmental stage and the number of endosperm nuclei. Endosperm nuclei counts were assigned an Endosperm Division Value (EDV), which estimates the number of divisions to reach the corresponding number of endosperm nuclei (Ungru et al., 2008). Mean EDV was calculated using the formula: 2^x^ = mean number of nuclei, where x is the EDV (x = LOG(mean number of nuclei)/LOG(2)).

Crosses with EE-GFP and TE1-GFP markers were imaged using the Andor DragonFly as described. Whole-mount imaging of seeds was performed using an Axioplan2 imaging microscope after 24 h/4°C incubation in 8:2:1 (w/v/v) chloral hydrate:water:glycerol (Grini et al., 2002). Mature dry seeds were imaged on a 1.5 x 2.5 cm grid using a Leica Z16apoA microscope connected to a Nikon D90 camera. Seed size and circularity were measured by converting images to black and white and then using the ImageJ “Threshold” and “Analyze particles” functions (https://imagej.nih.gov/ij/).

## 3 Results

### 3.1 Antagonistic effects on endosperm barriers in hybrid seeds from *A. lyrata and A. arenosa* crossed to *A. thaliana*

In order to compare the success of interspecific hybrids of *A. thaliana* crossed with *A. lyrata* or *A. arenosa*, we first observed seed-set. Hybrid crosses with *A. lyrata* fathers had significantly reduced seed set per silique compared to crosses with *A. arenosa* fathers or *A. thaliana* (Col-0) self crosses (Supplementary Figure S1A). The lower seed-set correlates with failure of pollen tube burst after entering the female gametophyte (Supplementary Figure S1B), a pre-zygotic barrier previously described (Escobar-Restrepo et al., 2007). Interestingly, pollen tube burst failure can also be observed in crosses between *A. thaliana* and *A. arenosa*, but to a lower degree, correlating with the observed seed-set frequency (Supplementary Figures S1A, B). Differences in seed set between the two hybrid crosses thus have a pre-zygotic base, and we conclude that a postzygotic hybridization barrier in these crosses is unaffected by the gametophyte induced reduction in seed-set.

*A. thaliana* self seeds and *A. thaliana* × *A. arenosa* or *A. thaliana* × *A. lyrata* hybrid seeds were comparable in size at 6 DAP (Figure 1A), but most *A. thaliana* × *A. lyrata* hybrid embryos were at an earlier developmental stage (Figure 1B). Large variation in embryonic stages including developmental arrest was observed in 12 DAP whole-mount chloral hydrate cleared hybrid seeds (Supplementary Figure S2A). A time series of Feulgen stained *A. thaliana* × *A. lyrata* hybrid seeds further identified variation in endosperm cellularization starting 3 DAP, resulting in developmental arrest at the globular embryo stage (Supplementary Figure S2B). The frequency of *A. thaliana* × *A. lyrata* early cellularization was higher than for *A. thaliana* self, and in strong contrast to the late cellularization in hybrid seeds from *A. thaliana* crossed with *A. arenosa* (Figure 1B; (Josefsson et al., 2006; Bjerkan et al., 2020)). The endosperm nuclei-number was also significantly lower in *A. thaliana* × *A. lyrata* compared to *A. thaliana* × *A. arenosa* hybrid seeds, suggesting a reduction in endosperm proliferation rate (Figure 1C). Nonetheless, germination rates of hybrid seeds from crosses with *A. lyrata* were significantly higher than crosses with *A. arenosa* (Figure 1D, Supplementary Datasheet S2), indicating that the endosperm hybrid barrier is influenced in a diametrically opposed manner, both in terms of the endosperm cellularization phenotype and the effective output measured as the ability of hybrid seeds to germinate.

**Figure 1:**
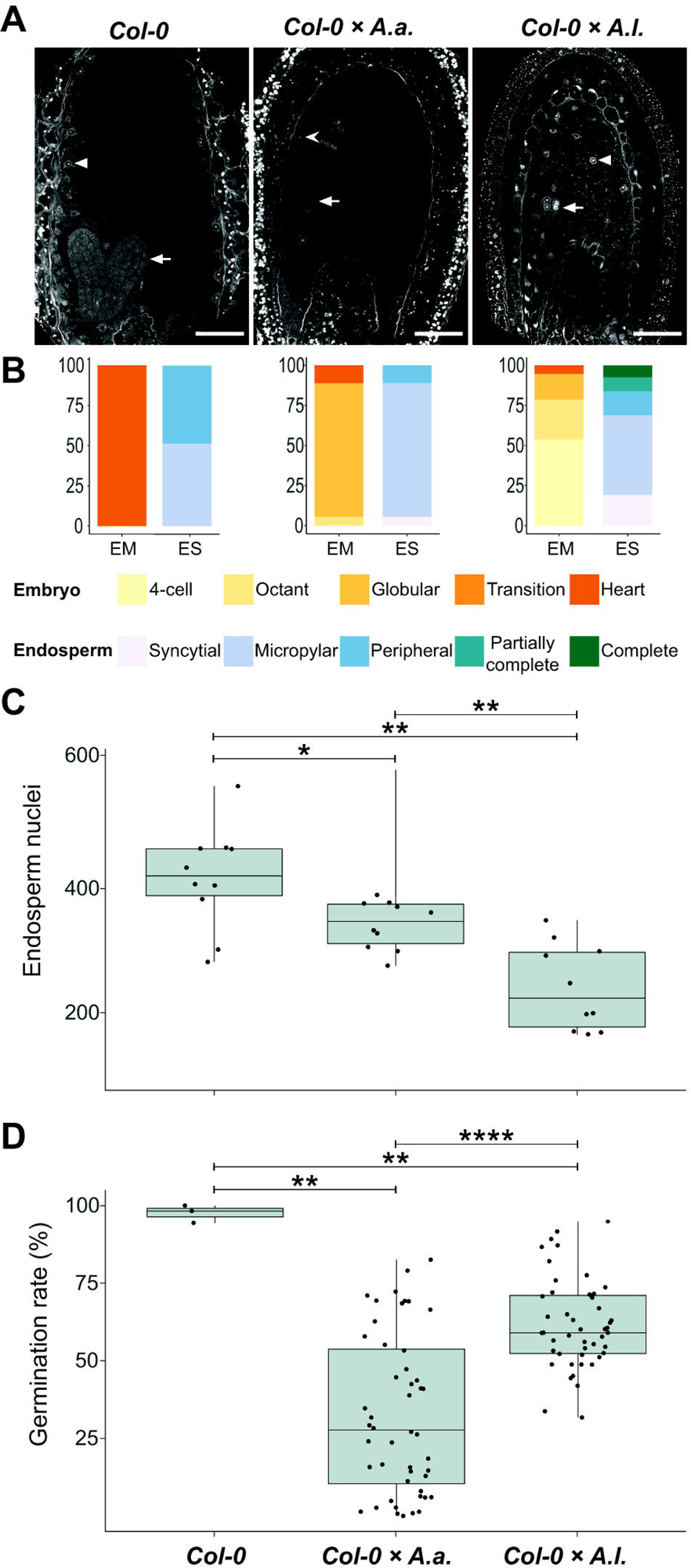
Antagonistic endosperm phenotypes in hybrid seeds. Seeds from crossing *A. arenosa* (*A.a.*) or *A. lyrata* (*A.l.*) as pollen donor to *A. thaliana* (Col-0) have antagonistic endosperm phenotypes and display opposing seed germination rates. **(a)** Confocal images of Feulgen-stained Col-0 and hybrid seeds 6 days after pollination (DAP) emphasizing endosperm cellularization. Open arrowhead points to syncytial endosperm nuclei, closed arrowheads point to cellularized endosperm nuclei and full arrows point to the embryo. In Col-0 self cellularization occurs at the 6 DAP embryo heart stage whereas in *A.a.* hybrid seeds the endosperm is mainly syncytial at 6 DAP. In *A.l.* hybrid seeds precocious endosperm cellularization is observed 6 DAP already at the early globular stage. All crosses are female × male. Scale bar = 50 μm. **(b)** Relative frequencies of embryo and endosperm stages in seeds from the same crosses as above. Col-0, n = 37; Col-0 × *A.a.*, n = 18; Col-0 × *A.l.*, n = 93; *EM*, embryo stages; ES, endosperm stages. **(c)** Number of endosperm nuclei in seeds from the same crosses as above, n = 10. **(d)** Germination rates in seeds from the same crosses as above. Biological replicates (siliques): Col-0, n = 4; Col-0 × *A.a.*, n = 48; Col-0 × *A.l.*, n = 48. Significance is indicated for the comparisons between all genotypes (Wilcoxon rank-sum test: \**P* ≤ 0.05; \*\**P* ≤ 0.01; \*\*\*\**P* ≤ 0.0001).

### 3.2 Endosperm temporal markers are shifted in hybrid seeds

Compared to *A. thaliana,* viable seeds resulting from crosses between *A. thaliana* mothers and *A. arenosa* or *A. lyrata* fathers appear to develop along a slower embryo and endosperm developmental path. This suggests that as long as endosperm cellularization and embryo development are synchronized, seed viability is not impacted. In *A. lyrata* × *A. arenosa* hybrid seeds with unsynchronized endosperm and embryo development, the embryo can be rescued *in vitro*, indicating that the barrier is caused by lack of nutrient support to the growing embryo (Lafon-Placette and Köhler, 2016).

To investigate synchronization of embryo and endosperm in *A. thaliana* (Col-0) hybrid seeds with *A. arenosa* or *A. lyrata* fathers, we used genetic markers of endosperm development that are expressed before and after endosperm cellularization in *A. thaliana (van Ekelenburg et al., 2022)*. In *A. thaliana* the EE-GFP genetic marker is expressed after fertilization and up to endosperm cellularization at 6-7 DAP (van Ekelenburg et al., 2022) and not expressed after cellularization (Figs 2, S3). In *A. thaliana* × *A. arenosa* hybrid seeds, expression of the EE-GFP marker was observed for the full duration of the time series (15 DAP) and no visible downregulation could be observed (Figs 2, S3). This expression pattern supports the observation that *A. thaliana* × *A. arenosa* hybrid seeds fail to initiate or have delayed endosperm cellularization (Figure 1) as described previously (Josefsson et al., 2006; Bjerkan et al., 2020). In contrast, when crossed to *A. lyrata*, expression of the EE-GFP marker decreased from 5 DAP and only a low frequency of seeds expressed the marker at cellularization around 9 DAP (Figs 2, S3). Taking the developmental delay in the *A. thaliana* × *A. lyrata* hybrid into account, the EE-GFP marker is prematurely terminated (Figure 2B), in accordance with the early cellularization phenotype (Figure 1). In *A. thaliana* self seeds, downregulation of the EE-GFP marker coincides with endosperm cellularization (van Ekelenburg et al., 2022). In hybrid seeds from *A. arenosa* or *A. lyrata* fathers, continued expression or premature EE-GFP downregulation, respectively (Figure 2B), coincides with diametrically opposed cellularization phenotypes both indicating that the endosperm phase change and cellularization is not synchronized with embryo development, leading to embryo and seed failure.

**Figure 2:**
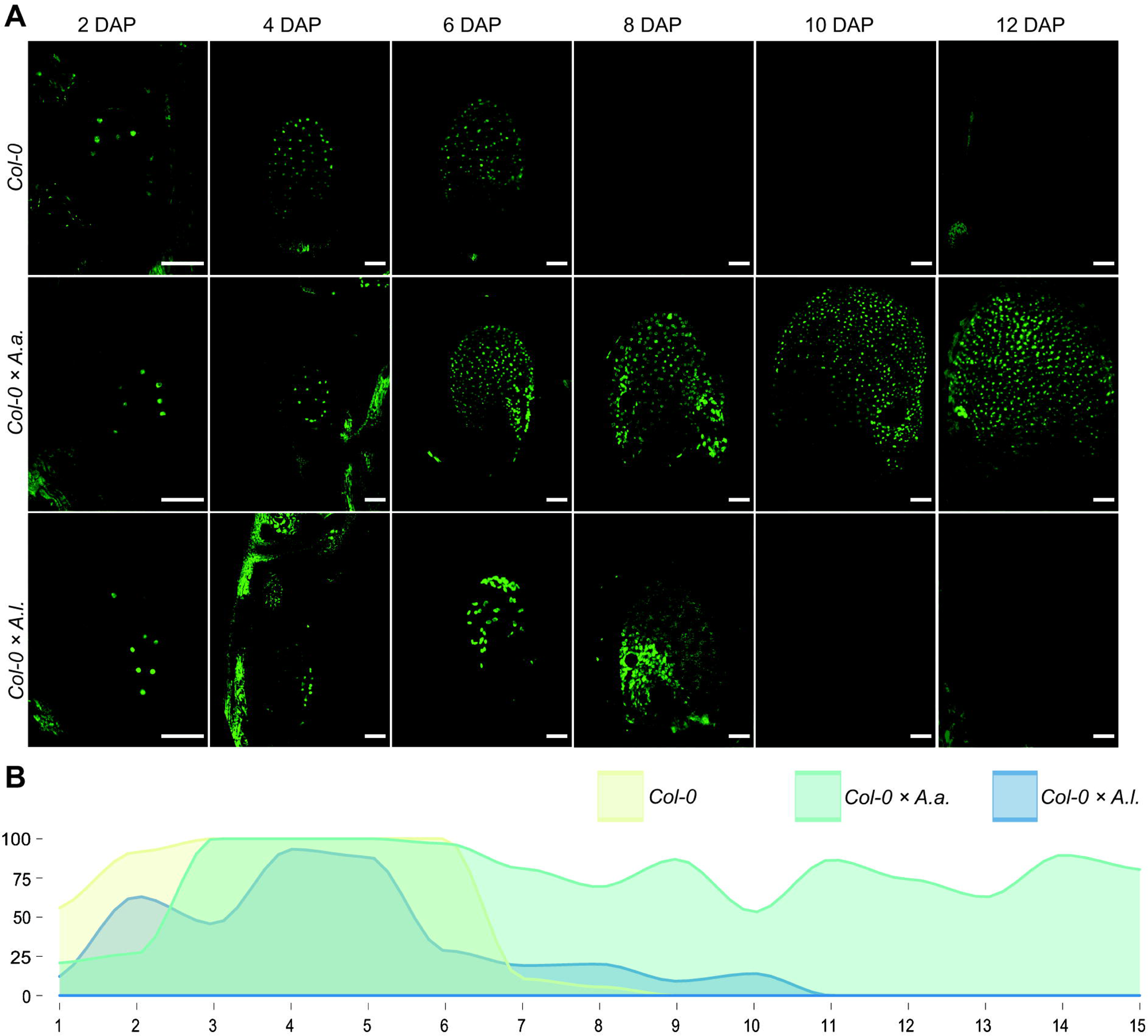
Early endosperm marker confirms aberrant cellularization timing in hybrid seeds. **(a)** Expression patterns of proAT5G09370>>H2A-GFP (EE-GFP) in seeds of *A. thaliana* (Col-0) and in hybrid seeds from crossing *A. arenosa* (*A.a.*) / *A. lyrata* (*A.l.*) as pollen donor to Col-0 at 2, 4, 6, 8, 10 and 12 days after pollination (DAP). Scale bar = 50 µm. **(b)** Percentage of hybrid seeds expressing EE-GFP at 1-15 DAP. *A. thaliana* (Col-0) self crosses and hybrid crosses are indicated with different colors.

In *A. thaliana,* the TE1-GFP genetic marker is expressed after cellularization at 6-7 DAP (van Ekelenburg et al., 2022) and until seed maturation at 17-19 DAP (Figs 3, S4). Delayed activation of marker expression was observed at low frequency in the *A. thaliana* × *A. arenosa* hybrid seeds with expression from 9 to 18 DAP (Figs 3, S4). Interestingly, in *A. thaliana* × *A. lyrata* hybrid seeds the TE1-GFP marker expressed prematurely from before 6 DAP globular stage seeds lasting until 18 DAP (Figs 3, S4), indicating premature endosperm phase change initiation and supporting the observed precocious endosperm cellularization phenotype (Figure 1).

**Figure 3:**
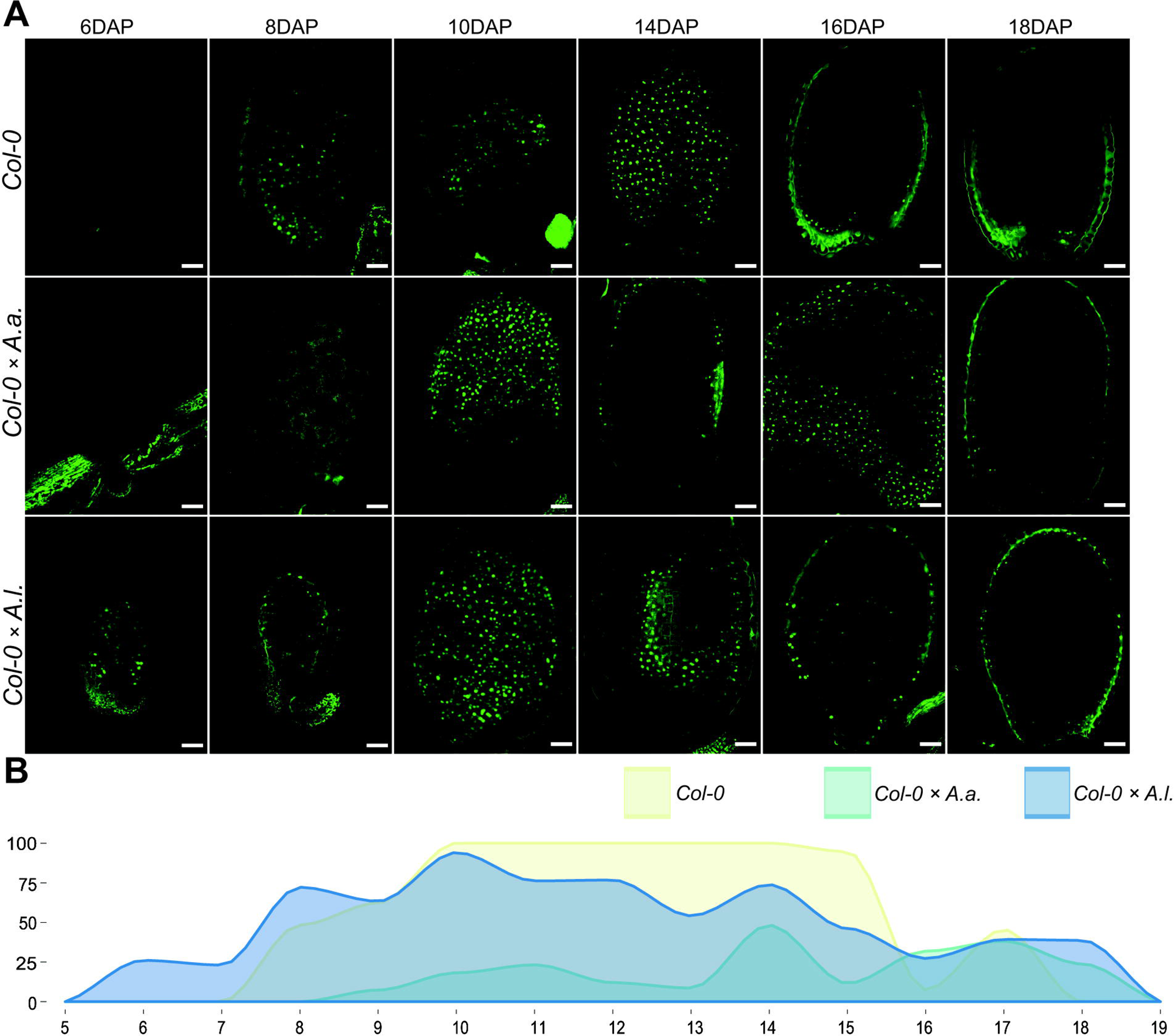
Total endosperm marker confirms aberrant cellularization timing in hybrid seeds. **(a)** Expression patterns of proAT4G00220>>H2A-GFP (TE1-GFP) in seeds of *A. thaliana* (Col-0) and in hybrid seeds from crossing *A. arenosa* (*A.a.*) / *A. lyrata* (*A.l.*) as pollen donor to *A. thaliana* (Col-0) at 6, 8, 10, 14, 16 and 18 days after pollination (DAP). Scale bar = 50 µm. **(b)** Percentage of hybrid seeds expressing TE1-GFP at 5-19 DAP. *A. thaliana* (Col-0) self crosses and hybrid crosses are indicated with different colors.

In *A. thaliana* seeds the decrease of EE-GFP expression and increase of TE1-GFP expression is strictly coordinated, with limited overlap. This is contrasted by the EE-GFP and TE1-GFP expression in seeds from hybrid crosses (Figure 4). Since the marker transgenes are expressed from the maternal *A. thaliana* genomes, the only difference in these crosses is the paternal contribution. For *A. thaliana* × *A. arenosa* an overlap in expression of the markers was observed from 9 DAP, caused by the prolonged expression of the EE-GFP marker (Figure 4). In *A. thaliana* × *A. lyrata* overlapping expression was observed from 6 to 10 DAP (Figure 4), showing a shift in TE1-GFP expression towards earlier developmental stages, although the *A. thaliana* × *A. lyrata* hybrid seeds develop slower compared to both *A. thaliana* selfed and *A. thaliana* × *A. arenosa* seeds (Figure 1).

**Figure 4:**
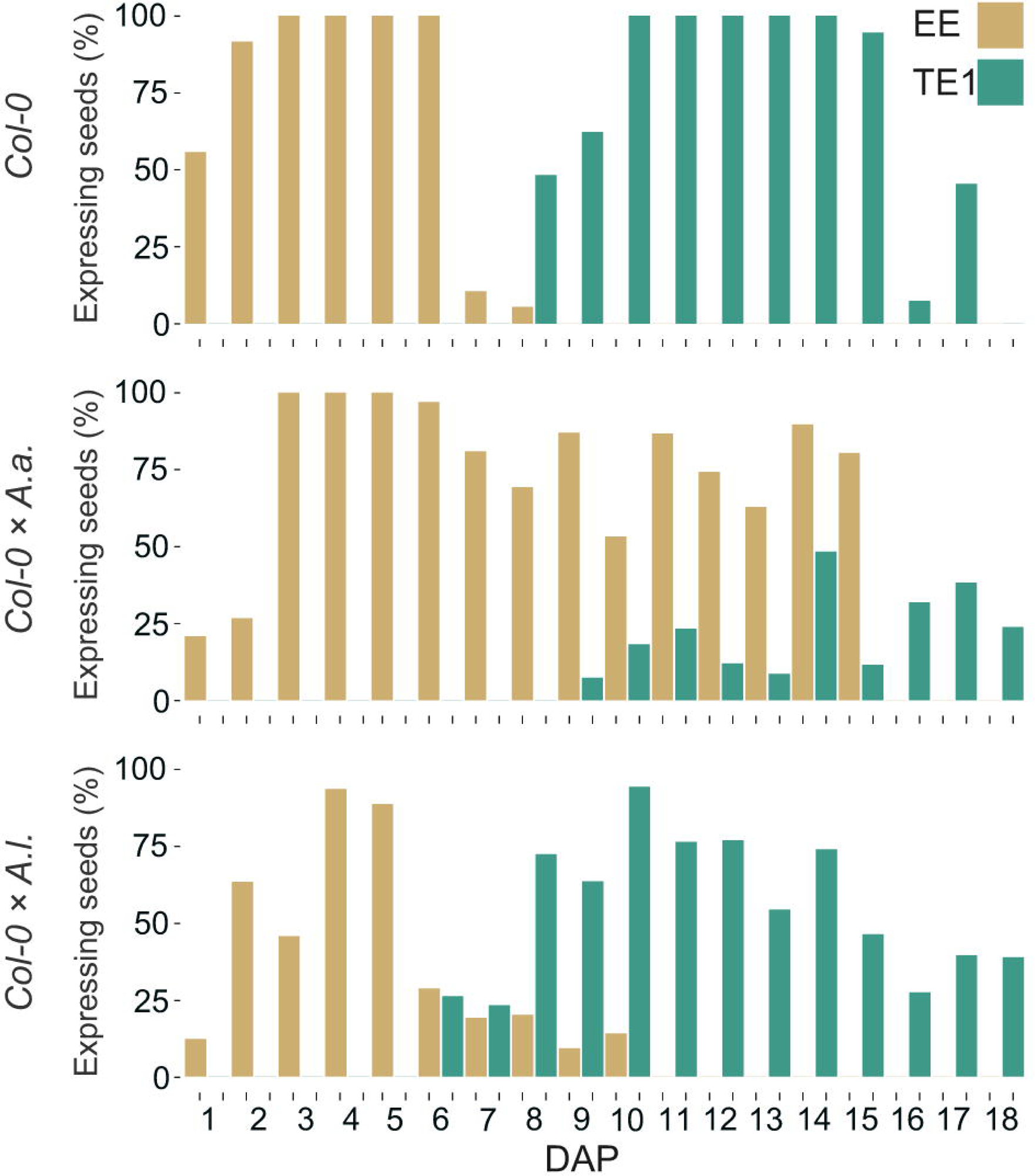
Overlapping expression of early and late endosperm markers in hybrid seeds. Percentage of seeds expressing proAT5G09370>>H2A-GFP (EE-GFP) and proAT4G00220>>H2A-GFP (TE1-GFP) at 1-18 days after pollination (DAP) in *A. thaliana* (Col-0) self-crosses and from crossing *A. arenosa* (*A.a.*) or *A. lyrata* (*A.l.*) as pollen donor to *A. thaliana* (Col-0). EE-GFP expression in the Col-0 × *A. arenosa* hybrid seeds were not documented after 15 DAP.

### 3.3 Temperature alters the hybrid barrier strength in diametrically opposed directions

Lowering of temperature from 22°C to 18°C ameliorates the germination efficiency of the hybrid seeds from *A. thaliana* mothers crossed to *A. arenosa* fathers (Bjerkan et al., 2020). In order to investigate if the phenotypically contrasting hybrid barrier observed in seeds from *A. lyrata* fathers was affected by temperature in a corresponding way, we performed crosses between *A. thaliana* (Col-0) mothers and *A. lyrata* or *A. arenosa* fathers at 4°C temperature windows ranging from 14°C to 26°C. Interestingly, the germination rate of *A. thaliana* × *A. arenosa* hybrid seeds were significantly enhanced by progressive lowering of the temperature, and contrasted by the germination rate of *A. thaliana* × *A. lyrata* hybrid seeds that was significantly enhanced by progressive increase of the temperature (Figure 5, Supplementary Datasheet S2). In the temperature window, the effects on the hybrid barriers followed a close to linear, but opposed reaction norm. We applied more extreme temperatures previously reported to be within the normal, and not stress inducing, growth-range of *A. thaliana* (Lloyd et al., 2018), but a further enhancement could not be obtained (Supplementary Figure S5).

**Figure 5:**
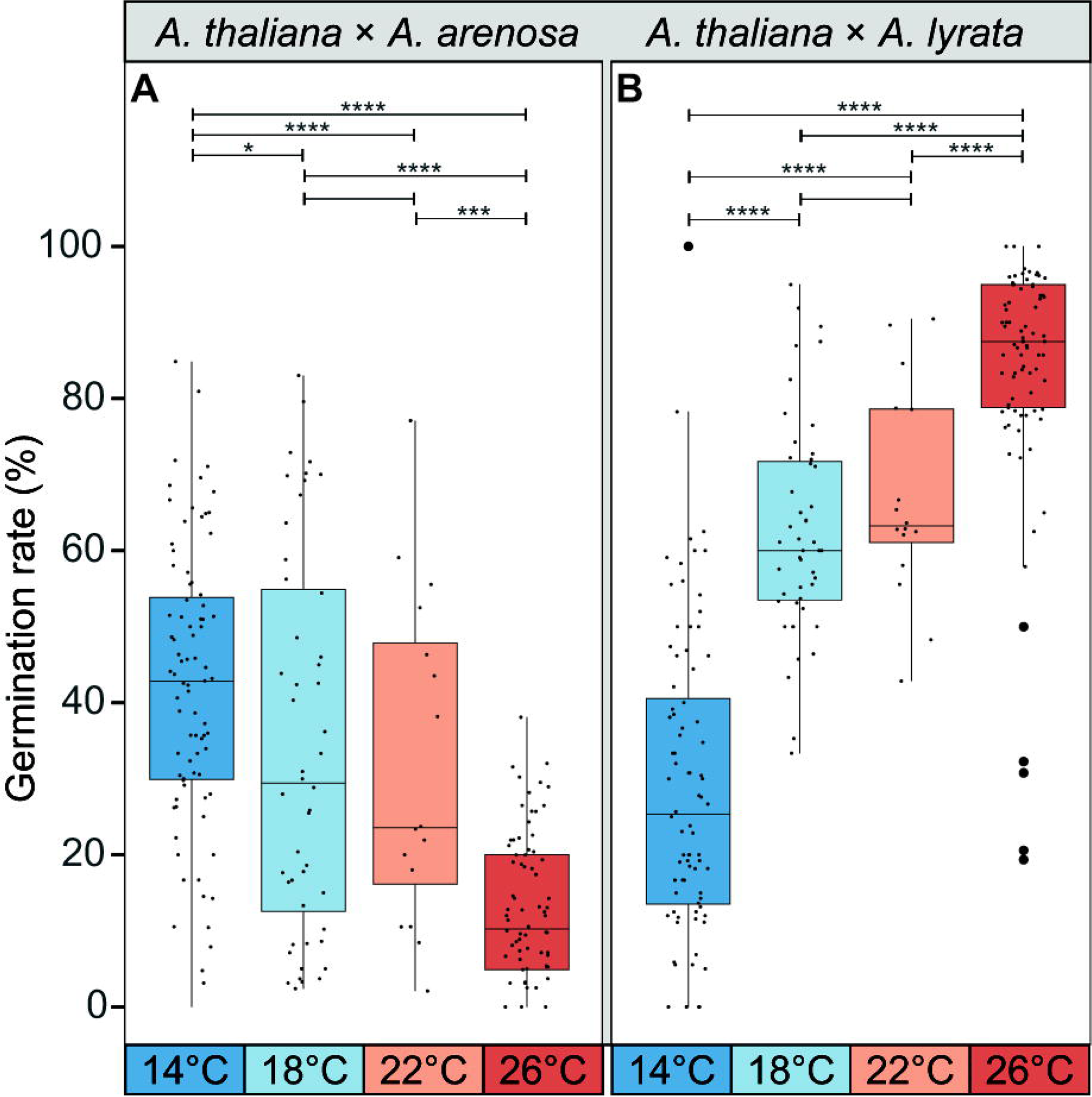
Temperature during seed development affects germination rate of hybrid seeds. **(a)** Germination rate of *A. thaliana* (Col-0) × *A. arenosa* hybrid seeds from crosses performed at 14°C, 18°C, 22°C and 26°C. Germination rate decreases with increasing temperature. Biological replicates (siliques): 14°C, n = 83 (3243 seeds); 18°C, n = 48 (2610 seeds); 22°C, n = 16 (796 seeds); 26°C, n = 78 (2713 seeds). **(b)** Germination rate of *A. thaliana* (Col-0) × *A. lyrata* hybrid seeds at 14°C, 18°C, 22°C and 26°C. Germination rate increases with increasing temperature. Biological replicates (siliques): 14°C, n = 84 (1676 seeds); 18°C, n = 47 (1540 seeds); 22°C, n = 16 (533 seeds); 26°C, n = 85 (2043 seeds). Box plot contains scattered data points representing germination rates observed per silique. Outliers are plotted as large data points. Significance is indicated for the comparisons between all temperatures (Wilcoxon rank-sum test: \**P* ≤ 0.05; \*\*\**P* ≤ 0.001; \*\*\*\**P* ≤ 0.0001).

### 3.4 Accessions of *A. thaliana* influence *A. arenosa* and *A. lyrata* hybrid barriers antagonistically

Different accessions of *A. thaliana* affect the strength of the hybrid barrier when crossed to *A. arenosa (Burkart-Waco et al., 2012; Bjerkan et al., 2020)*. Having demonstrated an antagonistic temperature effect on the hybrid barrier when *A. arenosa* or *A. lyrata* are crossed to *A. thaliana* mothers, we next investigated if different *A. thaliana* accessions influence the hybrid barrier in similar or opposite manner. We performed hybrid crosses with *A. arenosa* and *A. lyrata* fathers to the diploid *A. thaliana* accessions Col-0, C24 and Ws-2. Additionally, the tetraploid accession Wa-1 was used as a control, as tetraploid *A. thaliana* crossed to diploid *A. arenosa* has been shown to increase hybrid seed survival (Josefsson et al., 2006).

Compared to the Col-0 accession crosses, the C24 accession enhanced seed survival significantly when crossed to *A. arenosa*, contrasted by the *A. lyrata* hybrid where the germination rate declined highly significantly compared to the Col-0 cross (Figure 6; p ≤ 0.0001). For Ws-2, the germination rate was severely reduced in the *A. arenosa* hybrid cross, contrasted by moderately high (though lower than for Col-0) germination rate in the *A. lyrata* hybrid cross (Figure 6, Supplementary Datasheet S2).

**Figure 6:**
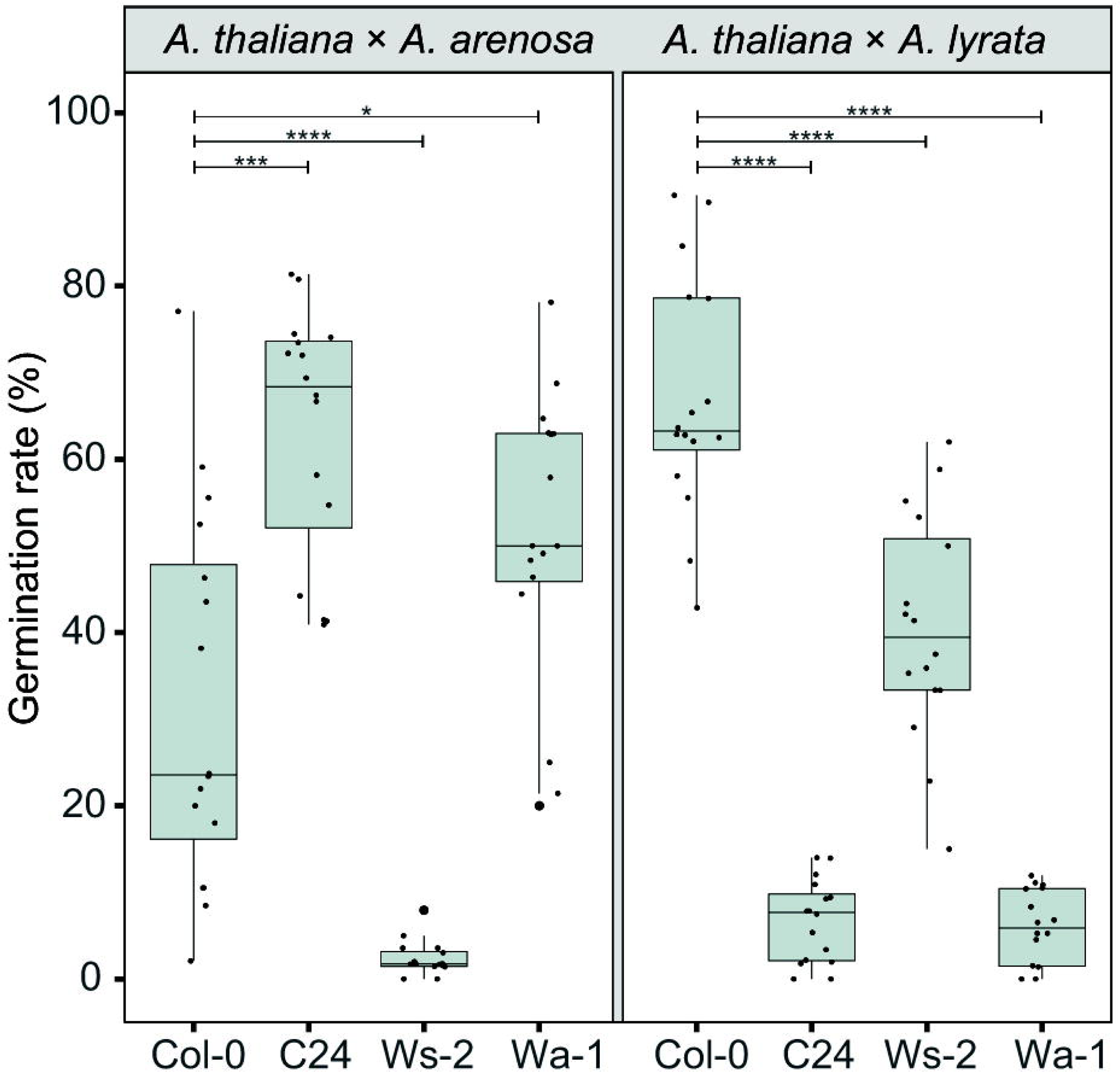
*A. thaliana* accessions affect hybrid barriers antagonistically. Germination rate of seeds from crossing *A. arenosa* (*A.a.*) / *A. lyrata* (*A.l.*) as pollen donor to *A. thaliana* (Col-0/C24/Ws-2/Wa-1) at 22°C. Biological replicates (siliques): Col-0 × *A.a.*, n = 16 (533 seeds); C24 × *A.a.*, n = 16 (805 seeds); Ws-2 × *A.a.*, n = 16 (947 seeds); Wa-1 × *A.a.*, n = 16 (805 seeds); Col-0 × *A.l.*, n = 16 (533 seeds); C24 × *A.l.*, n = 16 (821 seeds); Ws-2 × *A.l.*, n = 16 (572 seeds); Wa-1 × *A.l.*, n = 16 (759 seeds). Box plot contains scattered data points representing germination rates observed per silique. Outliers are plotted as large data points. Significance is indicated for comparisons between Col-0 × *A.a. /* Col-0 × *A.l.* and crosses involving other *A. thaliana* accessions (Wilcoxon rank-sum test; \**P* ≤ 0.05; \*\*\**P* ≤ 0.001; \*\*\*\**P* ≤ 0.0001).

In the tetraploid *A. thaliana* Wa-1 hybrid cross to diploid *A. arenosa*, the seed germination rate was enhanced, as previously reported (Josefsson et al., 2006). In contrast, in the Wa-1 hybrid cross to *A. lyrata*, the germination rate was significantly decreased compared to the Col-0 cross (Figure 6). Notably, the effect on seed viability was higher in the diploid accession C24 cross than with the tetraploid Wa-1 accession.

Crosses performed in parallel at 18°C and 22°C demonstrated the same trends at both temperatures (Supplementary Figure S6, Supplementary Datasheet S2). Although large differences in the barrier strength was observed, as measured by germination and seed viability, no obvious correlation could be found between seed survival and seed size and circularity (Supplementary Figure S7).

### 3.5 *A. thaliana* accession effects are not readily explained by endosperm cellularization phenotype

To investigate if the endosperm phenotype reflects the influence of accessions on hybrid seed viability, we inspected Feulgen stained 6 DAP hybrid seeds by confocal microscopy. We scored the number of endosperm nuclei in *A. thaliana* accessions and accession hybrids with *A. arenosa* and *A. lyrata* at 18°C and 22°C. The endosperm division value (EDV; (Ungru et al., 2008)) was generally higher when *A. arenosa* was involved (Figure 7, Supplementary Datasheet S3). No obvious correlation between the number of endosperm nuclei and hybrid seed viability was found (Supplementary Figure S8), however a significant correlation between endosperm proliferation rates and growth temperature could be observed in all crosses except *A. thaliana* C24 x *A. lyrata*. The latter hybrid cross did indeed exhibit very low germination rates (Figure 6), however similar low germination rates were found in *A. thaliana* Ws-2 x *A. arenosa* hybrid seeds but here accompanied by a high endosperm proliferation rate (Supplementary Figure S8, Supplementary Datasheet S3).

**Figure 7:**
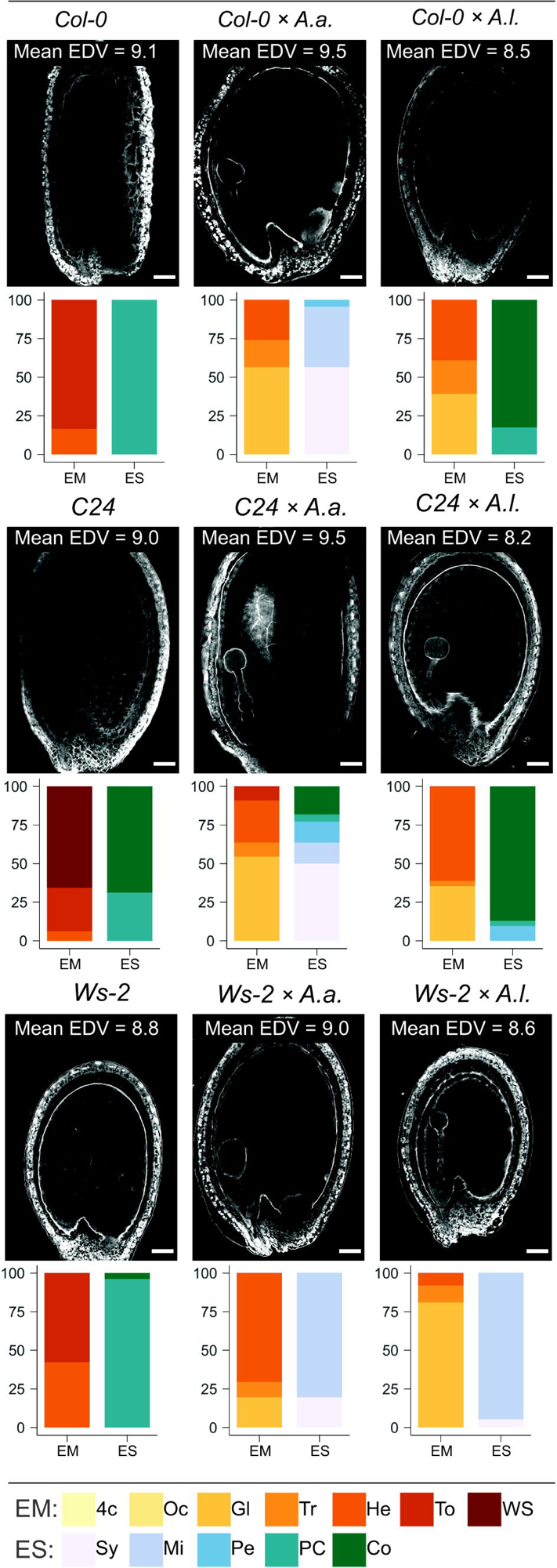
Effect of temperature, accession and hybridization on endosperm development. Confocal images showing endosperm cellularization of 6 DAP Feulgen-stained seeds from crossing *A. arenosa* (*A.a.*) or *A. lyrata* (*A.l.*) as pollen donor to *A. thaliana* (accession Col-0 or C24 or Ws-2) at 22°C. Scale bar = 50 μm. Mean endosperm division value (EDV) is shown within each image, n_EDV_ = 10 seeds. Beneath each image quantification of the described embryo and endosperm stages is shown as bar charts: Col-0, n = 36; Col-0 × A.a., n = 23; Col-0 × A.l., n = 23; C24, n = 32; C24 × *A.a.*, n = 22; C24 × *A.l.*, n = 31; WS-2, n = 26; WS-2 × *A.a.*, n = 51; WS-2 × *A.l.*, n = 37. Embryo stages (EM): 4c: 4-cell, Oc: Octant, Gl: Globular, Tr: Transition, He: Heart, To: Torpedo, WS: Walking stick. Endosperm cellularization stages (ES): Sy: Syncytial endosperm, Mi: Micropylar endosperm cellularization, Pe: Peripheral endosperm cellularization, PC: Partially complete endosperm cellularization, Co: Complete endosperm cellularization.

Seed phenotypes were scored for defined stages of embryo and endosperm development. *A. thaliana* accession self crosses at 22°C displayed embryo stages in the late heart to walking stick stage, with C24 exhibiting the fastest embryonic development. Endosperm cellularization was partly complete, though fully completed in most *A. thaliana* C24 self-seeds (Figure 7, Supplementary Datasheet S3). In *A. thaliana* accession x *A. arenosa* hybrid seeds, embryo development ranged from globular to transition stages. The endosperm was mainly syncytial or had initiated cellularization in the micropylar endosperm. In the C24 cross almost half of the seeds exhibited advanced cellularization stages and also complete cellularization. In this cross, higher germination rate was correlated with temporally correct timing of endosperm cellularization (Figs 6, 7).

In *A. thaliana* accession x *A. lyrata* hybrid seeds, embryonic stages ranged from globular to heart, where the C24 accession displayed a majority of heart stages, and Ws-2 a majority of globular stages. In the Col-0 and C24 accession hybrids, near uniform complete endosperm cellularization was observed, contrasted by early peripheral cellularization in Ws-2 (Figure 7). In the case of the latter hybrid cross, embryo development and endosperm cellularization appeared to be synchronized, leading to higher seed viability (Figure 6). However, large differences in seed viability between C24 (low) and Col-0 (high) hybrid crosses (Figure 6) was not reflected by the endosperm cellularization phenotype as both crosses had mostly fully cellularized endosperm and appeared to be in a similar embryonic stage (Figure 7). *A. thaliana* accession hybrid crosses at 18°C exhibited a similar pattern (Supplementary Figure S9). Major significant differences in seed viability between C24 and Ws-2 (low vs medium-high; Supplementary Figure S6) were not reflected by endosperm cellularization as both accession hybrids exhibited mostly micropylar endosperm (Supplementary Figure S9;). We conclude that the effect of using different accessions in the hybrid crosses can not readily be explained by a direct effect on the endosperm cellularization phenotype alone and that a more complex interaction between different genotypes occur.

### 3.6 Single gene mutation influences *A. arenosa* and *A. lyrata* hybrid barriers antagonistically

Deregulation of type I MADS-box TFs has been correlated with endosperm-based hybridization barriers (Walia et al., 2009), and many of these TFs are epigenetically regulated by the so-called FIS-PRC2 and the histone methyltransferase MEDEA (MEA) (Zhang et al., 2018). Mutation of *MEA* results in ectopic proliferation of endosperm nuclei and delayed cellularization (Grossniklaus et al., 1998; Köhler et al., 2003; Guitton et al., 2004) and we therefore investigated if endosperm overproliferation in *A. thaliana mea* mutant mothers crossed to *A. arenosa* or *A. lyrata* enhances or alleviates the hybrid barriers, respectively.

Heterozygous self-crossed *A. thaliana mea* mutants resulted in a reduced germination rate of 60% meaning that 80% of seeds carrying the mutant maternal allele failed to germinate due to delayed endosperm cellularization (Supplementary Figure S10). Crossing *A. thaliana mea* to *A. arenosa* or *A. lyrata* resulted in a significant decrease in seed survival compared to Col-0 crosses (Supplementary Figure S10, Supplementary Datasheet S2). Reduced germination rate in both crosses corresponded to an additive effect of the reduced germination of the heterozygous *A. thaliana mea* mutation. Compared to expected values, single gene mutation of *mea* could not bypass the *A. thaliana* x *A. lyrata* species barrier, nor enhance the *A. thaliana* x *A. arenosa* barrier (Supplementary Figure S10, Supplementary Datasheet S2).

We studied the effect of single candidate genes regulated by FIS-PRC2 (Zhang et al., 2018). The mutant *agl35-1* in the Col-0 background was previously shown to strengthen the barrier when *A. thaliana* was crossed to *A. arenosa* (Bjerkan et al., 2020) and *AGL35* was upregulated in the same hybrid cross (Walia et al., 2009). *AGL40* is similarly expressed in the endosperm (Zhang et al., 2018), upregulated in hybrids (Walia et al., 2009) and mutant seeds have reduced seed size (Kirkbride et al., 2019). To investigate if mutation of single candidate genes could produce opposed effects on the hybrid barrier when crossing *A. thaliana* mothers to *A. lyrata* or *A. arenosa*, as observed when changing temperature or *A. thaliana* accession (Figs 5, 6), we performed *A. lyrata* and *A. arenosa* crosses to *A. thaliana agl35-1* and *agl40-1* and scored seed germination.

Interestingly, in crosses where *agl35-1* was crossed to *A. arenosa* or *A. lyrata* a highly significant decrease or increase in germination rate was observed, respectively, compared to wild type Col-0 crosses (Figure 8A). Single mutation of *AGL35* affected the hybrid barrier strength in diametrically opposed directions, as in crosses to *A. lyrata* the germination rate was significantly enhanced, in contrast to *A. thaliana* x *A. arenosa* crosses where the germination rate was significantly reduced (Figure 8A). Mutation of *AGL40* crossed to *A. arenosa* did not significantly affect germination rate, but in crosses to *A. lyrata* germination rate was significantly reduced. Col-0 crossed to *A. lyrata* displayed reduced seed size (Figs S2, S7) due to early endosperm cellularization (Figure 1), and thus the mutation of *AGL40* may increase the frequency of early cellularization.

**Figure 8:**
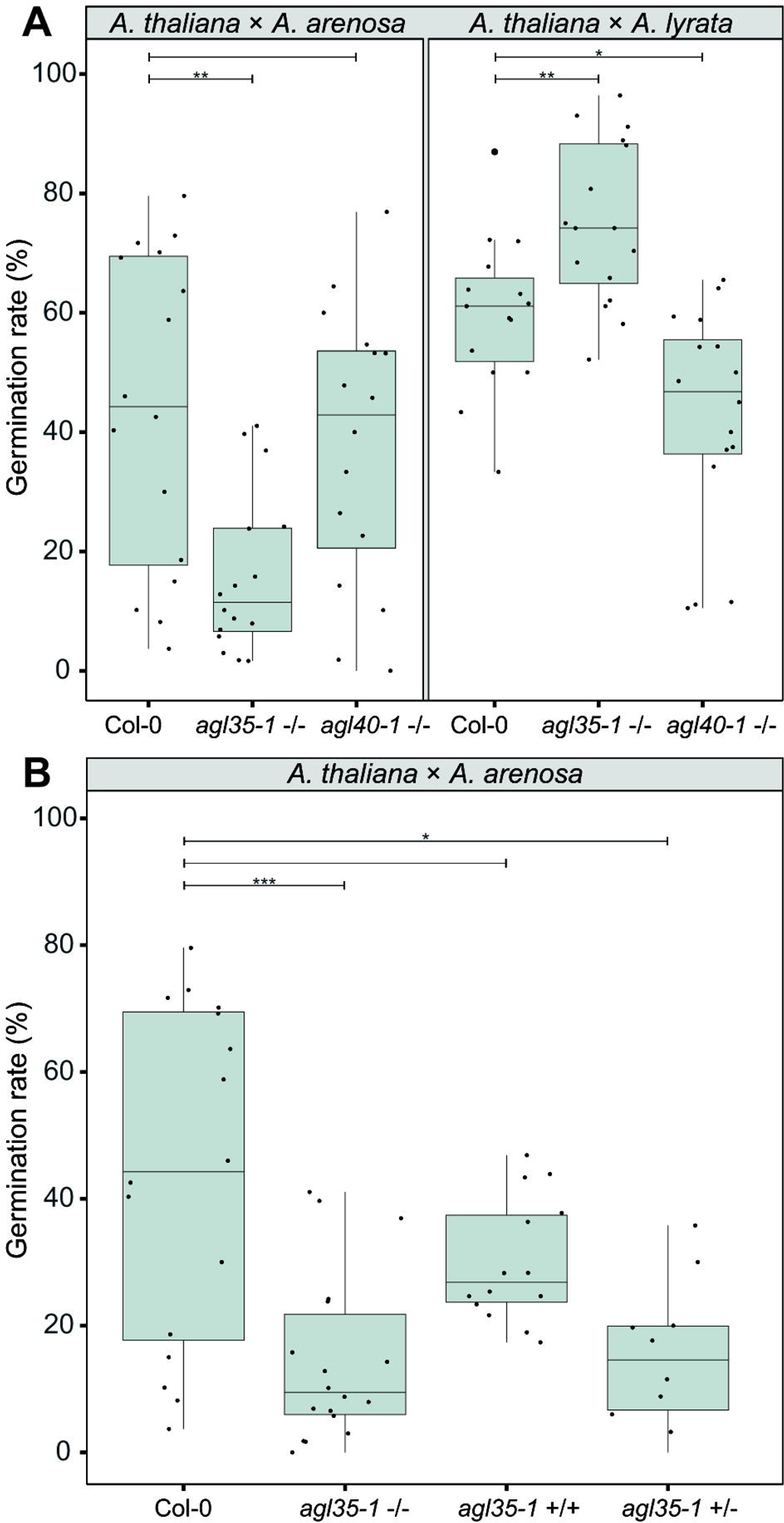
Genetic parameters influence the hybrid barrier. **(a)** Germination rate of seeds from crossing *A. arenosa* (*A.a.*) / *A. lyrata* (*A.l.*) as pollen donors to *A. thaliana* (Col-0), single mutants *agl35-1*-/- and *agl40-1*-/- at 18°C. Biological replicates (siliques): Col-0 × *A.a.*, n = 16 (833 seeds); *agl35-1*-/- × *A.a.*, n = 16 (904 seeds); *agl40-1*-/- × *A.a.*, n = 16 (879 seeds); Col-0 × *A.l.*, n = 16 (457 seeds); *agl35-1*-/- × *A.l.*, n = 16 (509 seeds); *agl40-1*-/- × *A.l.*, n = 16 (525 seeds). **(b)** Germination rate of seeds from crossing *A. arenosa* (*A.a.*) as pollen donor to *A. thaliana* (Col-0), homozygous *agl35-1*-/-, twice introgressed *agl35*-/- into Col-0 (*agl35*+/+), and heterozygous *agl35-1 +/-* at 18°C. Biological replicates (siliques): Col-0 × *A.a.*, n = 16 (833 seeds); *agl35-1*-/- × *A.a.*, n = 18 (1064 seeds); *agl35*+/+ × *A.a.*, n = 14 (1562 seeds); *agl35-1 +/-* × *A.a.*, n = 10 (525 seeds). Box plot contains scattered data points representing germination rates observed per silique. Outliers are plotted as large data points. Significance is indicated for the comparisons between Col-0 crosses and the mutant crosses (Wilcoxon rank-sum test; *P ≤ 0.05; **P ≤ 0.01; ***P ≤ 0.001).

To test if cryptic genetic variation in the *agl35-1* mutant line could account for the observed phenotype, heterozygous *agl35-1* was introgressed twice to Col-0 and segregating progeny of selfed heterozygotes were crossed to *A. arenosa.* The segregating *agl35-1 A. thaliana* mothers were genotyped for the *agl35-1* insert and germination rate of hybrid seeds was scored (Figure 8B). Importantly, the germination rate of segregating wildtype plants (*agl35+/+)* was not significantly different from wildtype (Col-0) when crossed to *A. arenosa.* Both homozygous (*agl35-1 -/-)* and heterozygous *agl35-1 +/-* plants crossed to *A. arenosa* displayed significantly lower germination rate than Col-0 crossed to *A. arenosa*, indicating that the increased strength of the hybrid barrier was caused by mutation of *AGL35*. Furthermore, the observation that the strength of the hybrid barrier can be caused by heterozygous (*agl35-1 +/-*) plants crossed to *A. arenosa* indicates that the observed phenotype is caused by genetic interaction occurring in the fertilization products, the embryo or the endosperm.

## 4 Discussion

Interspecies hybrid seeds from crossing *A. lyrata* or *A. arenosa* as the paternal parent to *A. thaliana* mothers show antagonistic endosperm cellularization phenotypes, with late cellularization in crosses with *A. arenosa* and early cellularization in crosses with *A. lyrata*. In both cases, cellularization failure results in an endosperm-based hybrid barrier and reduced viability of germinating seeds. This compares to previous studies where timing of endosperm cellularization is influenced by the paternal species in reciprocal *A. arenosa* and *A. lyrata* interspecies crosses and crosses within the genus *Capsella* (Rebernig et al., 2015; Lafon-Placette et al., 2017). Intriguingly, we find that a temperature gradient leads to diametrically opposed cellularization phenotype responses in hybrid endosperm with *A. arenosa* or *A. lyrata* as pollen donors. In addition, *A. thaliana* accession genotypes also influence hybrid seed viability in opposite directions. To this end, we demonstrate that single gene mutation in *A. thaliana* MADS-box TF *AGL35* independently can affect the germination rates of *A. arenosa* or *A. lyrata* hybrid seeds in opposite directions.

### 4.1 Ectopic timing of endosperm developmental phase change

The endosperm genetic markers EE-GFP and TE1-GFP mark syncytial endosperm development before cellularization and cellular endosperm stages, respectively (van Ekelenburg et al., 2023). We show that timing of the developmental phase change connected to endosperm cellularization is disturbed in hybrid seeds. In the Col-0 × *A. arenosa* hybrid seeds that fail to cellularize, the EE-GFP marker continues to be expressed throughout endosperm development, indicating phase change failure. In the Col-0 × *A. lyrata* hybrid seeds, characterized by early cellularization, the developmental time point of TE1-GFP expression indicates occurrence of a premature phase change. These findings support that not only the timing of endosperm cellularization is affected in these developing hybrid seeds, but also the developmental timing of the genetic network associated with endosperm phase change and maturation occurring at cellularization. In accordance with the incomplete endosperm hybridization barrier, prolonged expression of EE-GFP in Col-0 × *A. arenosa* seeds and precocious expression of TE1-GFP in Col-0 × *A. lyrata* seeds were not observed in all individual seeds. This suggests that gene regulation associated with the endosperm phase change within each hybrid seed varies and potentially is affected by genetic or epigenetic variation that modulate threshold levels for gene activation or repression. Paternal excess in *A. thaliana* inter-ploidy crosses results in failure of endosperm cellularization (Scott et al., 1998; Hehenberger et al., 2012; Martinez et al., 2018) and a recent report demonstrates that inter-ploidy uncellularized endosperm was correlated with increased abscisic acid (ABA) levels, suggesting that endosperm cellularization is connected to dehydration responses in the developing embryo (Xu et al., 2023).

In our system the sole difference between the two hybrids is the paternal parent, indicating a trans-acting mechanism, where differential expression from *A. lyrata* and *A. arenosa* genomes regulates the genetic markers expressed from the *A. thaliana* genome. Supporting this hypothesis, paternal transmission of mutants in NRPD1 (NUCLEAR RNA POLYMERASE D1) can bypass the cellularization phenotype in paternal excess inter-ploidy crosses (Erdmann et al., 2017; Martinez et al., 2018). NRPD1 is a main component in the the RNA-directed DNA methylation (RdDM) pathway, resulting in small RNA directed gene regulation by *de novo* DNA methylation (Law et al., 2013; Kirkbride et al., 2019) and could be a potential trans-acting regulatory mechanism (Erdmann et al., 2017). Future experiments to identify transcriptional differences from the parental genomes in hybrid seeds may point at key genes and mechanisms responsible for ectopic timing of the endosperm developmental phase change.

### 4.2 Temperature and accession affect the viability of hybrid seeds

We found that by increasing the temperature from 14°C to 26°C, Col-0 × *A. arenosa* seeds display significantly decreased germination rates (more than 30%). The same temperature range has an opposite effect in Col-0 × *A. lyrata* seeds resulting in increased germination rates (more than 50%). This demonstrates a temperature dependent genetic mechanism that acts antagonistically when *A. thaliana* is crossed to *A. arenosa* or *A. lyrata* and produces diametrically opposed cellularization phenotype responses in the hybrid endosperm.

The current knowledge on the effect of temperature in early seed development is limited (Paul et al., 2020), but an effect of temperature on hybrid seed development in reciprocal crosses of wheat and barley has previously been reported (Molnár-Láng and Sutka, 1994). The sensitivity of endosperm cellularization to heat stress during early endosperm development has been demonstrated in rice (Folsom et al., 2014) and type I MADS-box TFs are deregulated during moderate heat stress (Chen et al., 2016). Interestingly, in *Brassica oleracea*, temperature affects ABA levels specifically in the endosperm and cooler temperatures obstruct the breakdown of ABA in the desiccating endosperm (Chen et al., 2021). This is consistent with the aforementioned finding that inter-ploidy uncellularized endosperm correlated with increased abscisic acid (ABA) levels, suggesting that endosperm cellularization is connected to dehydration responses in the developing embryo (Xu et al., 2023). ABA catabolism in response to temperature may therefore be a potent mechanism to explain the temperature influence on the hybrid barrier when *A. thaliana* is crossed to *A. arenosa*. In a similar manner, we speculate that precocious cellularization in crosses with *A. lyrata* may be associated with a similar mechanism that triggers ABA breakdown, but this needs further investigation.

Notably, the effect of using different *A. thaliana* accessions in the hybrid crosses is larger than the temperature effect (close to 70% difference), and even larger than the interploidy effect. While the Col-0 and Ws-2 *A. thaliana* accessions resulted in a generally higher germination rate when hybridized with *A. lyrata* compared to *A. arenosa*, the C24 accession had the opposite effect. The way the accessions affected hybrid seed viability in opposite directions may point at a similar mechanism as observed in the temperature experiment. However, our results do not readily explain the observed germination rates by cellularization phenotype alone, and further investigations are required to resolve these observations.

Crossing tetraploid Wa-1 *A. thaliana* accession to diploid *A. arenosa* increases hybrid seed survival (Josefsson et al., 2006). In addition, ploidy affects the strength of the hybrid barrier in crosses between *A. arenosa* and *A. lyrata*, where higher ploidy in *A. lyrata* increases the hybrid seed survival rate, while higher ploidy in *A. arenosa* causes total seed lethality (Lafon-Placette et al., 2017). Our data show that a similar effect is found using the diploid accession C24, suggesting that C24 may have a higher effective ploidy and Endosperm Balance Number (EBN; (Johnston and Hanneman, 1982)) compared to Col-0 and Ws-2. This corresponds well with the hypothesis that *A. lyrata* has a lower EBN compared to *A. arenosa* (Lafon-Placette and Köhler, 2016), explaining why crosses with C24 or Wa-1 decrease seed viability in the *A. lyrata* hybrid. However, it does not explain why *A. lyrata* crosses with the diploid C24 is more detrimental than crosses with the tetraploid Wa-1, suggesting that accession genotypes, in addition to ploidy, has an effect on the endosperm-based hybridization barrier.

### 4.3 *AGL35* influences endosperm cellularization in hybrid seeds

AGAMOUS-LIKE (AGL) type I MADS-box TFs are highly expressed in the seed, specifically during endosperm cellularization (Bemer et al., 2010; Zhang et al., 2018; Bjerkan et al., 2020). Their importance in the endosperm-based hybridization barrier has been suggested by several studies (Josefsson et al., 2006; Walia et al., 2009; Bjerkan et al., 2020) and it has been hypothesized that timing of endosperm cellularization requires a stoichiometric balance between members of different MADS-box protein complexes (Batista et al., 2019). In this study we demonstrate that mutation in *A. thaliana AGL35* has a highly significant and opposite effect on the hybrid barrier phenotype when crossed to *A. lyrata* and *A. arenosa*, respectively. AGL35 is bi-allelicly expressed in the chalazal endosperm (Bemer et al., 2010; Bjerkan et al., 2020) and upregulated in crosses between *A. thaliana* and *A. arenosa* compared with compatible crosses (Walia et al., 2009). Our results indicate that *AGL35* is involved in the transition from syncytial to a cellularized endosperm, and may function as a promoter of cellularization, as mutant crosses to *A. arenosa* result in lower seed survival, while mutant crosses to *A. lyrata* results in increased survival compared to Col-0 crosses. Interestingly, a massive-multiplexed yeast two-hybrid study identified interaction between AGL62 and AGL35 (Trigg et al., 2017). These AGL TFs have seemingly antagonistic functions as AGL62 is a suppressor of endosperm cellularization (Kang et al., 2008). Paternal excess interploidy crosses cause increased AGL62 expression, correlated with endosperm cellularization failure (Erilova et al., 2009). AGL62 is also a direct target of the FIS PRC2 complex (Hehenberger et al., 2012) whereas we could see no direct effect on the *A. arenosa* or *A. lyrata* hybrid with *A. thaliana* by mutation of FIS PRC2. The antagonistic effects of single gene mutation of *AGL35* is intriguing, and we speculate that expression differences between *A. arenosa* and *A. lyrata* in the hybrid endosperm may account for our observations but future investigation of this interaction and the role of AGL35 in regulation of endosperm-based hybridization barriers is required.

### 4.4 Conclusions

The findings in this study introduce a rigorous model system for the dissection of the influence of abiotic and genetic parameters in hybrid admixture, and have a large potential to support breeding and climate research. Further examination and usage of these approaches could help pinpoint genes, networks or gene dosage balances that are involved in overcoming the endosperm-based hybridization barrier. Species previously thought to be unable to hybridize due to postzygotic seed lethality may be able to do so given favorable conditions, and a similar effect could also apply to interploidy hybrids.

Currently, it is not known if the temperature effect on hybridization success is mediated by the same genetic network that is operated by changes in ploidy or genetic variation. Phenotypically, the temperature effect restores defects in timing of cellularization, but it is not known if the trigger is upstream or downstream of the causative genetic network. Elucidation of the genetic, epigenetic and mechanistic basis for this cross talk between the genic and environmental factors is therefore essential for our understanding of the plasticity of endosperm-based hybridization barriers.

## Supporting information

Datasheet S1 - S4

## Acknowledgment

We thank the Laboratory of Flow Cytometry, Institute of Botany, Academy of Sciences (Czech Republic) for assistance.

## Competing interests

The authors state no competing interests.

## Author contributions

PEG, RMA, KNB & AKB designed the research; RMA, IVM & KNB performed the experiments; PEG, RMA, KNB & AKB analyzed and discussed the data; PEG, RMA, KNB & AKB wrote the article; All authors revised and approved the article.

## Data availability

Raw data and R-scripts can be found at https://github.com/PaulGrini/ArabidopsisEndospermBarriers

## Supplementary material

**Supplementary Figure S1:** Reduced seed set and pollen tube burst failure in *A. thaliana* (Col-0) × *A. lyrata* (*A.l.*) F1 hybrids.

**Supplementary Figure S2:** Seed phenotypes in F1 hybrids.

**Supplementary Figure S3:** Confocal micrographs of proAT5G09370>>H2A-GFP (EE-GFP) in seeds of *A. thaliana* (Col-0) and in hybrid seeds.

**Supplementary Figure S4:** Confocal micrographs of proAT4G00220>>H2A-GFP (TE1-GFP) in seeds of *A. thaliana* (Col-0) and in hybrid seeds.

**Supplementary Figure S5:** Temperature affects germination rate of hybrid seeds. **Supplementary Figure S6:** Temperature effect on germination rate in hybrid seeds varies with *A. thaliana* accession.

**Supplementary Figure S7:** Effect of temperature, *A. thaliana* accession and hybridization on seed size and circularity.

**Supplementary Figure S8:** Number of endosperm nuclei varies with temperature, accession and hybridization.

**Supplementary Figure S9:** Effect of temperature, accession and hybridization on endosperm development.

**Supplementary Figure S10:** Germination rate of *A. thaliana mea* and WT hybrid seeds from crosses with *A. arenosa* or *A. lyrata*.

**Supplementary Datasheet S1**: Flow Cytometry

**Supplementary Datasheet S2**: Germination assays

**Supplementary Datasheet S3**: Number of nuclei in crosses

**Supplementary Datasheet S4**: Feulgen phenotype observations

## Notes

https://github.com/PaulGrini/ArabidopsisEndospermBarriers

### Competing Interest Statement

The authors have declared no competing interest.

