## Supplemental Figures S1-S10 for "Genetic and environmental manipulation of *Arabidopsis* hybridization barriers uncover antagonistic functions in endosperm cellularization"

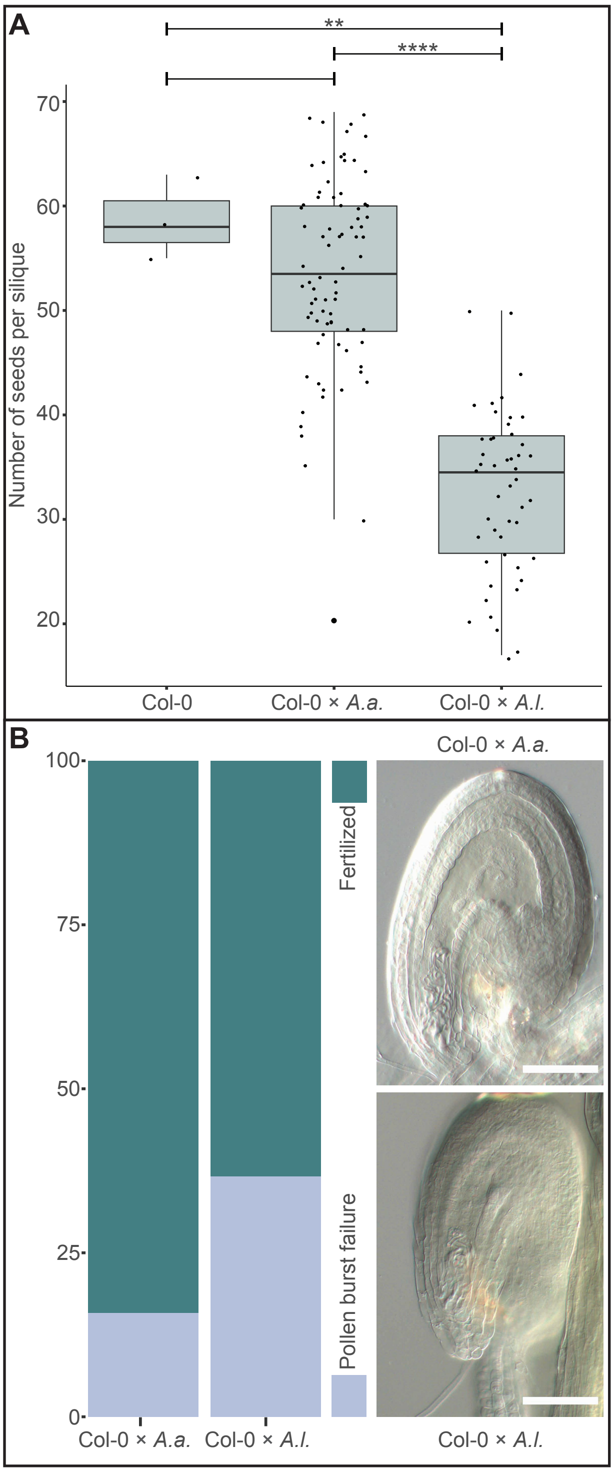


**Supplementary Figure 1.** Reduced seed set and pollen tube burst failure in *A. thaliana* (Col-0) × *A. lyrata* (*A.l.*) F1 hybrids. (A) Number of seeds per silique. Biological replicates (siliques): Col-0, n = 3; Col-0 × *A. arenosa* (*A.a.*), n = 78; Col-0 × *A.l.*, n = 48. Outliers are plotted as large data points. Significance is indicated for comparisons between Col-0 and Col-0 × A.a./A.l. (Wilcoxon rank-sum test: **P ≤ 0.01, ****P ≤ 0.0001). (B) Left: Percentage of seeds with pollen tube burst failure in Col-0 × *A.a.* (n = 57 seeds) and Col-0 × *A.l.* (n = 202 seeds) growing at 22°C. Right: Light microscopy images of chloral hydrate cleared seeds at 3 days after pollination (DAP) from Col-0 × *A.l.*/*A.a.* showing pollen tube burst failure. Scale bar = 50 µm.

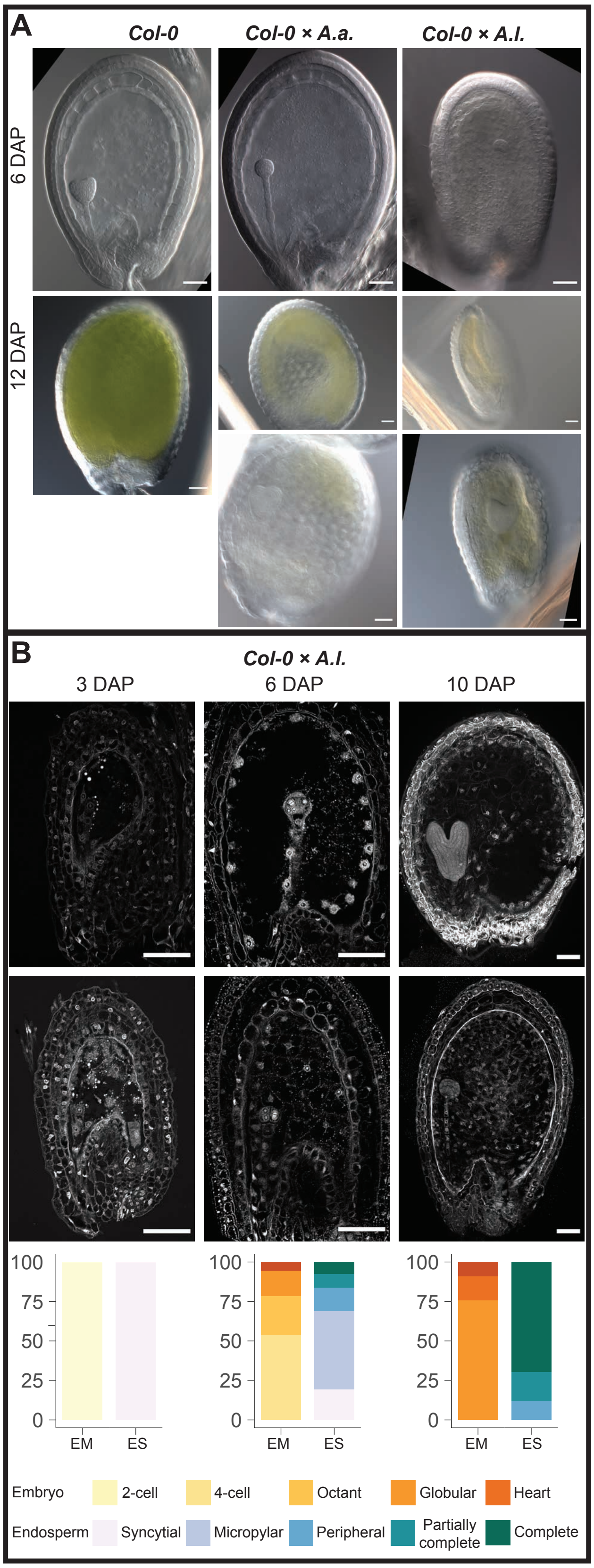


**Supplementary Figure 2.** Seed phenotypes in F1 hybrids. (A) Light microscopy images of chloral hydrate cleared seeds at 6 and 12 days after pollination (DAP) from A. thaliana (Col-0), Col-0 × A. arenosa (A.a.) and Col-0 × A. lyrata (A.l.). () Confocal micrographs showing endosperm cellularization of Feulgen-stained seeds at 3, 6 and 10 DAP from Col-0 × A.l. Frequency of endosperm cellularization and embryo stages is indicated by color code. Biological replicates (siliques): 3 DAP, n = 72; 6 DAP, n = 93; 10 DAP, n = 33; EM (embryo stages); ES (endosperm stages). Scale bar = 50 µm.

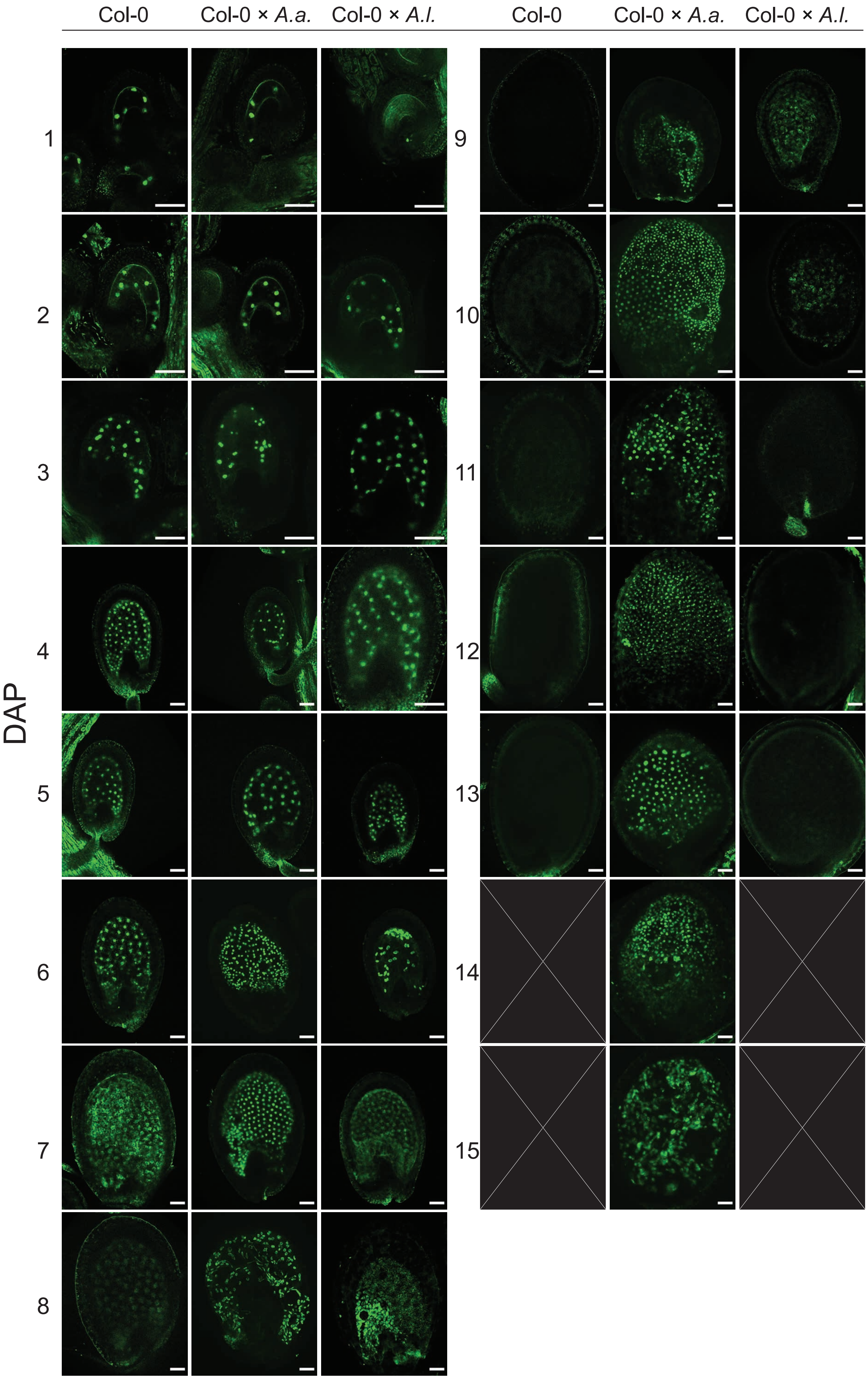


**Supplementary Figure 3.** Confocal micrographs of proAT5G09370>>H2A-GFP (EE-GFP) in seeds of *A. thaliana* (Col-0) and in hybrid seeds. Crosses of *A. thaliana* (Col-0) with *A. arenosa* (*A.a.*) or *A. lyrata* (*A.l.*) used as pollen donors from 1 to 15 days after pollination (DAP) are shown. Scale bar = 50 µm.

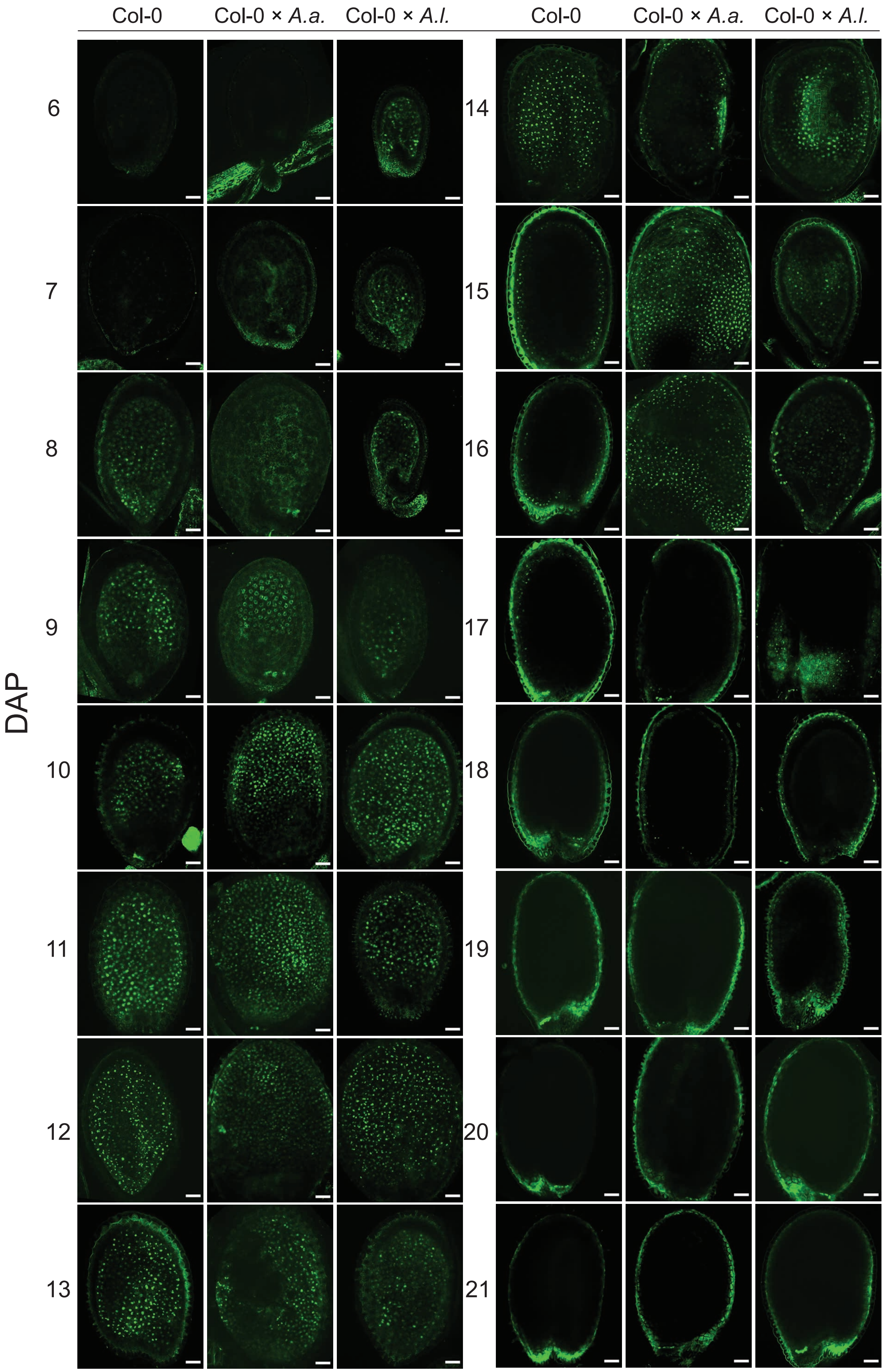


**Supplementary Figure 4.** Confocal micrographs of proAT4G00220>>H2A-GFP (TE1-GFP) in seeds of *A. thaliana* (Col-0) and in hybrid seeds. Crosses of *A. thaliana* (Col-0) with *A. arenosa* (*A.a.*) or *A. lyrata* (*A.l.*) used as pollen donors from 6 to 21 days after pollination (DAP) are shown. Scale bar = 50 µm.

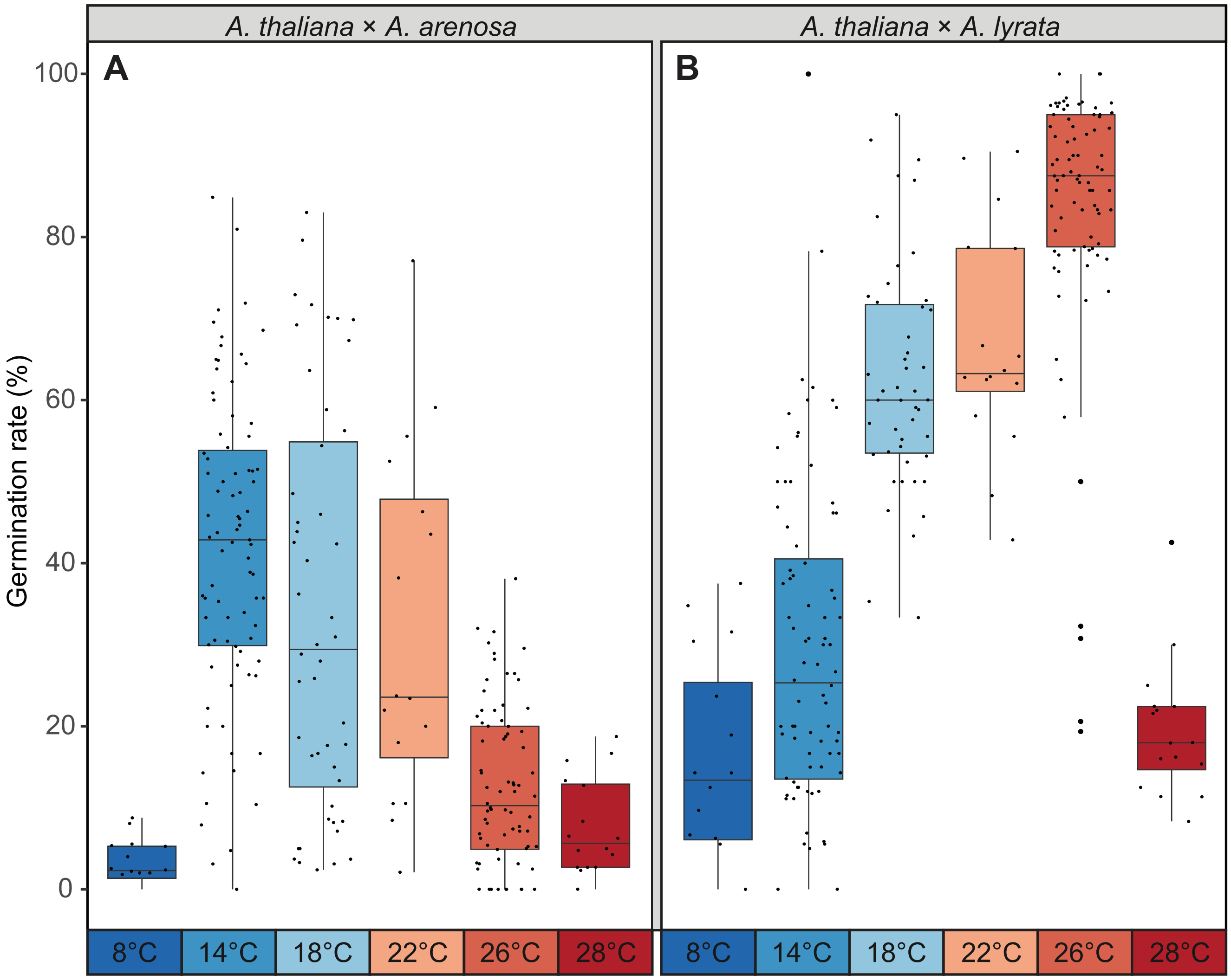


**Supplementary Figure 5.** Temperature affects germination rate of hybrid seeds. (A) Germination rate of *A. thaliana* (Col-0) × *A. arenosa* (*A.a.*) hybrid seeds at 8°C, 14°C, 18°C, 22°C, 26°C and 28°C. Biological replicates (siliques): 8°C, n = 16 (811 seeds); 14°C, n = 83 (3243 seeds); 18°C, n = 48 (2610 seeds); 22°C, n = 16 (796 seeds); 26°C, n = 78 (2713 seeds); 28°C, n = 16 (663 seeds). (B) Germination rate of *A. thaliana* (Col-0) × *A. lyrata* (*A.l.*) hybrid seeds at 8°C, 14°C, 18°C, 22°C, 26°C and 28°C. Biological replicates (siliques): 8°C, n = 16 (345 seeds); 14°C, n = 84 (1676 seeds); 18°C, n = 47 (1540 seeds); 22°C, n = 16 (533 seeds); 26°C, n = 85 (2043 seeds); 28°C, n = 16 (699 seeds). Box plot contains scattered data points representing germination rates observed per silique. Outliers are plotted as large data points.

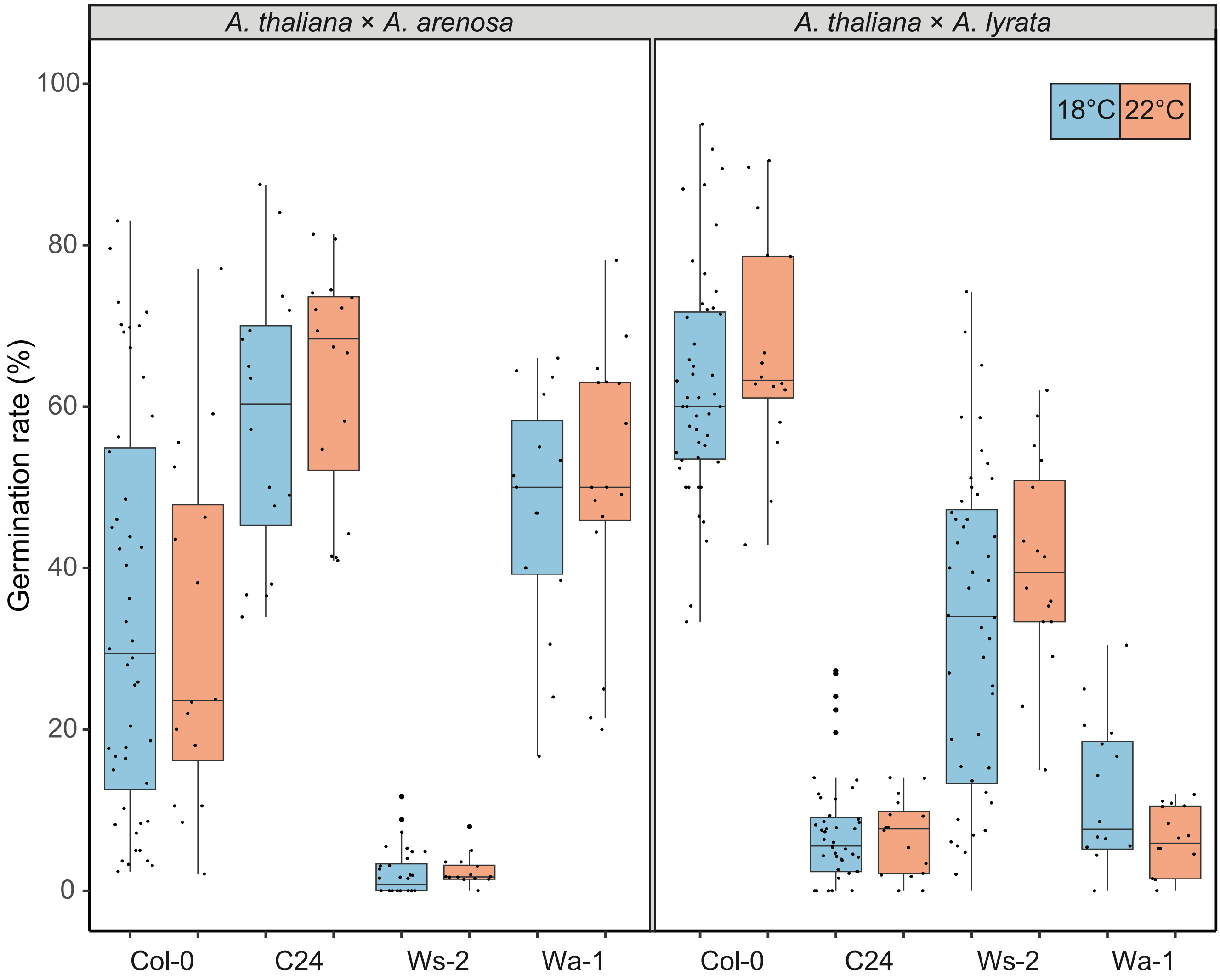


**Supplementary Figure 6.** Temperature effect on germination rate in hybrid seeds varies with *A. thaliana* accession. Germination rate of seeds from crossing *A. arenosa* (*A.a.*) / *A. lyrata* (*A.l.*) as pollen donor to *A. thaliana* (Col-0/C24/Ws-2/Wa-1) at 18°C and 22°C. Biological replicates (siliques): Col-0 × *A.a.* 18°C, n = 48 (2610 seeds); Col-0 × A.a. 22°C, n = 16 (533 seeds); C24 × *A.a.* 18°C, n = 16 (925 seeds); C24 × *A.a.* 22°C, n = 16 (805 seeds); Ws-2 × *A.a.* 18°C, n = 16 (987 seeds); Ws-2 × *A.a.* 22°C, n = 16 (947 seeds); Wa-1 × *A.a.* 18°C, n = 15 (546 seeds); Wa-1 × *A.a.* 22°C, n = 16 (805 seeds); Col-0 × *A.l.* 18°C, n = 47 (1540 seeds); Col-0 × *A.l.* 22°C, n = 16 (533 seeds); C24 × *A.l.* 18°C, n = 16 (801 seeds); C24 × *A.l.* 22°C, n = 16 (821 seeds); Ws-2 × *A.l.* 18°C, n = 16 (843 seeds); Ws-2 × *A.l.* 22°C, n = 16 (572 seeds); Wa-1 × *A.l.* 18°C, n = 16 (546 seeds); Wa-1 × *A.l.* 22°C, n = 16 (759 seeds). Box plot contains scattered data points representing germination rates observed per silique. Outliers are plotted as large data points.

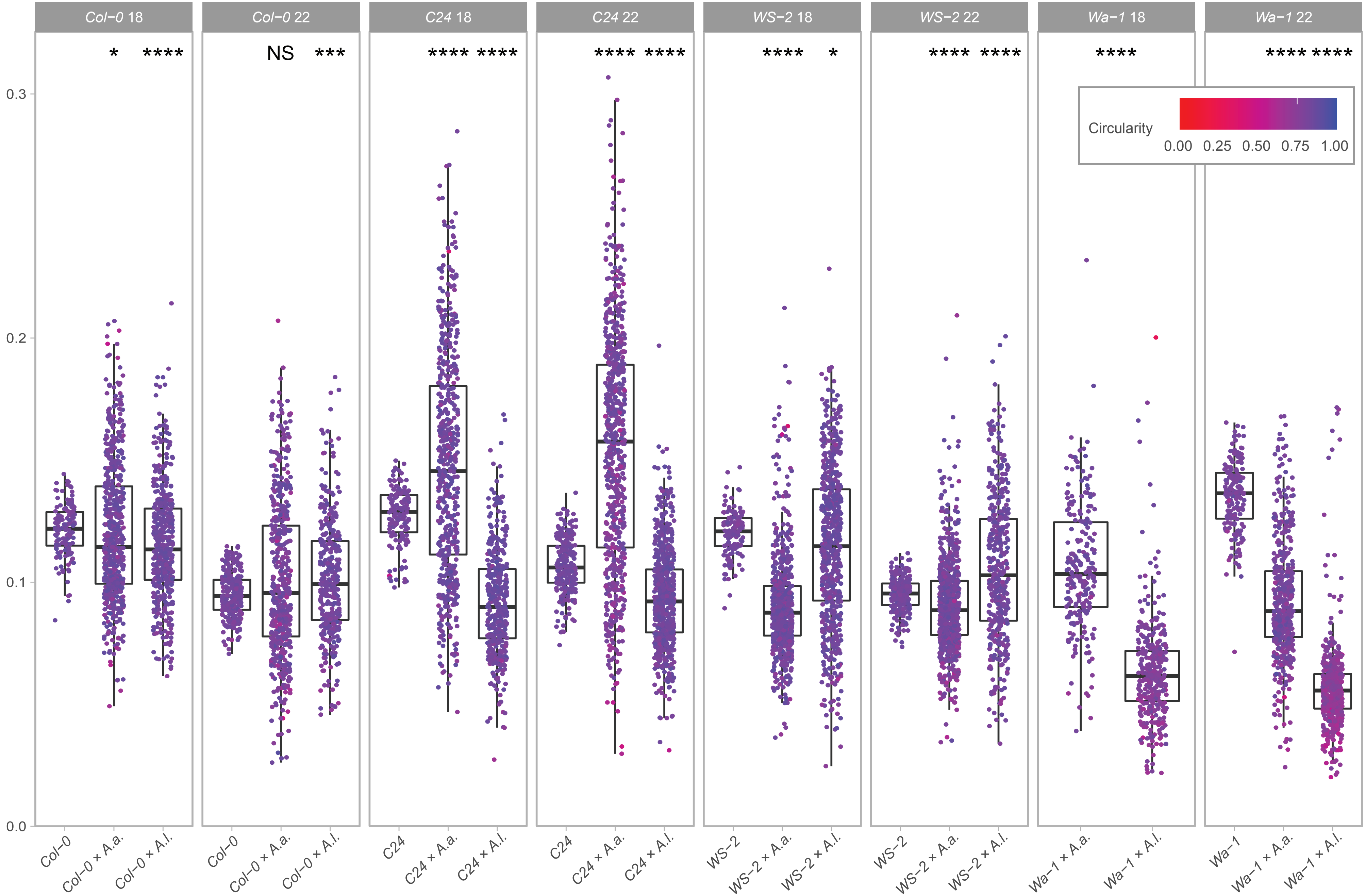


**Supplementary Figure 7.** Effect of temperature, *A. thaliana* accession and hybridization on seed size and circularity. Boxplot of seed size with circularity shown in a color gradient from 0 (low degree of circularity) to 1 (high degree of circularity) from A. thaliana (Col-0/C24/WS-2/Wa-1) self-crosses and from crossing *A. arenosa* (*A.a.*) / *A. lyrata* (*A.l.*) as pollen donor to Col-0/C24/WS-2/Wa-1. Col-0 18°C, n = 147; Col-0 x *A.a.* 18°C, n = 590; Col-0 x *A.l.* 18°C, n = 483; Col-0 22°C, n = 278; Col-0 x *A.a.* 22°C, n = 524; Col-0 x *A.l.* 22°C, n = 343; C24 18°C, n = 153; C24 x *A.a.* 18°C, n = 571; C24 x *A.l.* 18°C, n = 399; C24 22°C, n = 237; C24 x *A.a.* 22°C, n = 711; C24 x *A.l.* 22°C, n = 671; WS-2 18°C, n = 125; WS-2 x *A.a.* 18°C, n = 640; WS-2 x *A.l.* 18°C, n = 652; WS-2 22°C, n = 236; WS-2 x *A.a.* 22°C, n = 768; WS-2 x *A.l.* 22°C, n = 405. Significant differences in seed size are indicated for the comparisons between *A. thaliana* self-cross and accession crosses for each accession and temperature, except for Wa-1 at 18°C where the comparison was between Wa-1 × *A.a.* and Wa-1 × *A.l.* (Wilcoxon rank-sum test: NS P > 0.05; *P ≤ 0.05; ***P ≤ 0.001); ****P ≤ 0.0001.

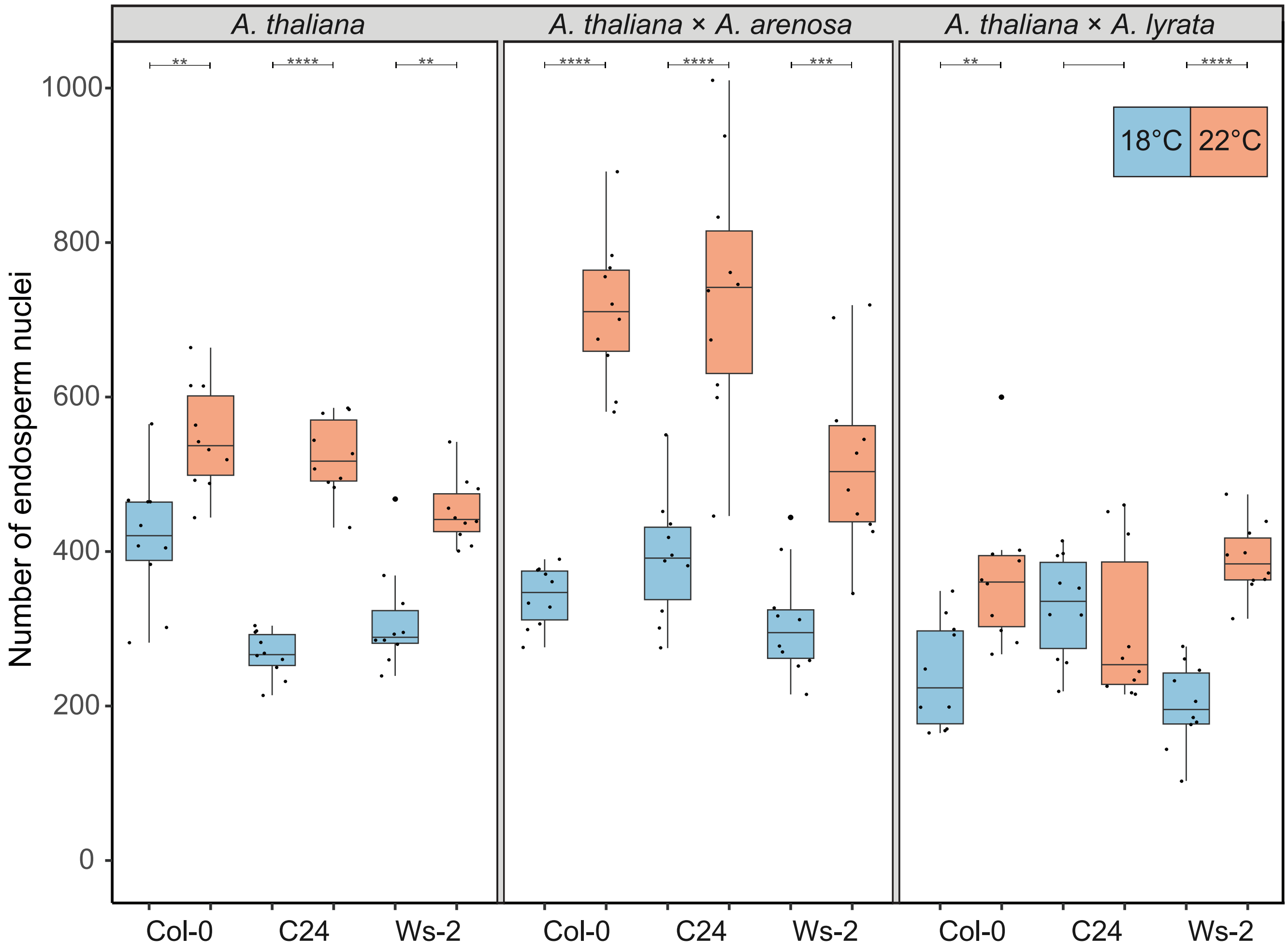


**Supplementary Figure 8.** Number of endosperm nuclei varies with temperature, accession and hybridization. Number of endosperm nuclei was counted in seeds (n = 10) from *A. thaliana* (Col-0/C24/Ws-2) self-crosses and from crossing *A. arenosa* (*A.a.*) or *A. lyrata* (*A.l.*) as pollen donor to *A. thaliana* (Col-0/C24/Ws-2). Significance is indicated for the comparisons of 18°C and 22°C between all accessions (Wilcoxon rank-sum test: $**P \leq 0.01$; $***P \leq 0.001$; $****P \leq 0.0001$). Outliers are plotted as large data points.

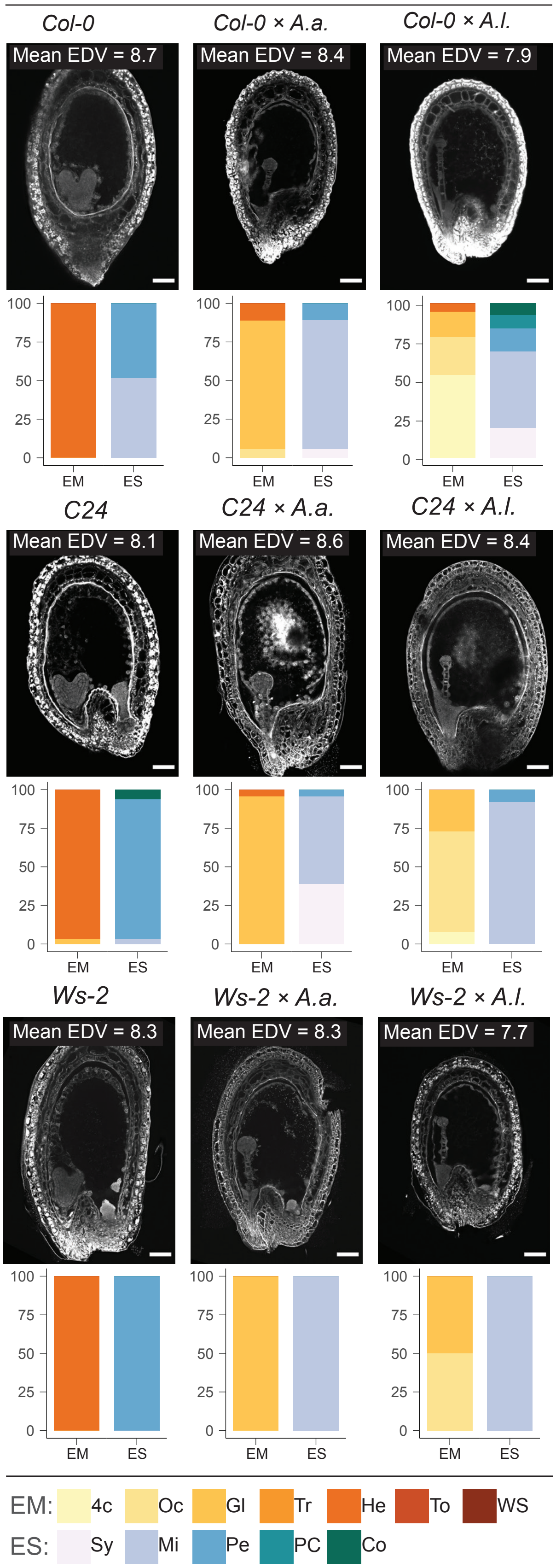


**Supplementary Figure 9.** Effect of temperature, accession and hybridization on endosperm development. Confocal images showing endosperm cellularization of Feulgen-stained seeds from *A. arenosa* (*A.a.*) or *A. lyrata* (*A.l.*) crossed as pollen donor to *A. thaliana* (accession Col-0 or C24 or Ws-2) at 18°C. Scale bar = 50 µm. Mean endosperm division value (EDV) is shown within each image, nEDV = 10 seeds. Quantification of the described embryo and endosperm stages in each cross are shown as bar charts: Col-0, n = 37; Col-0 × *A.a.*, n = 18; Col-0 × *A.l.*, n = 93; C24, n = 32; C24 × *A.a.*, n = 23; C24 × *A.l.*, n = 37; WS-2, n = 23; WS-2 × *A.a.*, n = 16; WS-2 × *A.l.*, n = 12. Embryo stages (EM): 4c: 4-cell, Oc: Octant, Gl: Globular, Tr: Transition, He: Heart, To: Torpedo, WS: Walking stick. Endosperm cellularization stages (ES): Sy: Syncytial, Mi: Micropylar, Pe: Peripheral, PC: Partially complete, Co: Complete.

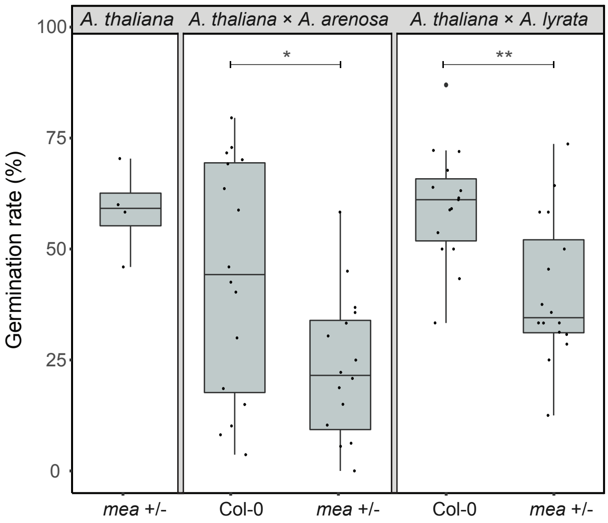


**Supplementary Figure 10.** Germination rate of *A. thaliana mea* and WT hybrid seeds from crosses with *A. arenosa* or *A. lyrata*. The x-axis indicates the genotype of the *A. thaliana* cross partner. Crossing *mea* to *A. arenosa* or *A. lyrata* results in a significant decrease in seed survival when compared to Col-0 crossed to A. arenosa or *A. lyrata*. The *mea* mutant crossed to self results in 60% viable seeds, indicating that the crosses to *A. arenosa* and *A. lyrata* do not have an impact on the seed survival compared to the Col-0 background. Wilcoxon rank-sum test: $^*P \leq 0.05$; $^{**}P \leq 0.01$. Outliers are plotted as large data points.
